# Inference of host-pathogen interaction matrices from genome-wide polymorphism data

**DOI:** 10.1101/2023.07.06.547816

**Authors:** Hanna Märkle, Sona John, Lukas Metzger, STOP-HCV Consortium, M Azim Ansari, Vincent Pedergnana, Aurélien Tellier

## Abstract

Coevolution is defined as the evolutionary change in one antagonist (host) in response to changes in the other antagonist (pathogen). At the genetic level, these changes are determined by genotype x genotype (GxG) interactions. We build on a general theoretical model of a host-pathogen interaction to derive four indices to retrieve key features of GxG interactions. The four developed indices extract relevant information from polymorphism data of randomly sampled uninfected hosts as well as infected hosts and their respective pathogen strains. Using these indices as summary statistics in an Approximate Bayesian Computation method, we can show their power to discriminate between GxG interaction matrices. Second, we apply our ABC method to a SNP data set of 451 European humans and their infecting Hepatitis C Virus (HCV) strains supplemented by polymorphism data of 503 individuals from the 1,000 genomes project. As our indices encompass and extend previous natural co-GWAs we recover many of the associations previously reported for this dataset and infer their underlying interaction matrix. We reveal a new candidate gene for resistance to HCV in the human genome, and two groups of significant GxG associations exhibiting gene-for-gene interactions. We suggest that the inferred types of GxG interactions result from the recent expansion, adaptation and low prevalence of the HCV virus population in Europe.

**Significance statement:** Why are some host individuals susceptible/resistant to infection by certain pathogen genotypes and others not? Understanding the genetic characteristics of genes driving host-pathogen interactions is crucial to predict epidemics. We develop four indices based on a mathematical model and build a Bayesian statistical method computing these indices on full genome data of infected hosts and their infecting pathogen strains and data of non-infected hosts. We can pinpoint the genes underlying host-pathogen interactions and infer their characteristics. Applying our framework to data from European humans and the Hepatitis C virus, we discover a new potential resistance gene in humans and reveal how the virus has adapted in the last 150 years to match the genetic diversity of the European human population.

## Introduction

Host-pathogen or host-parasite antagonistic interactions are pervasive in nature. Their relevance ranges from specific simple interactions underpinning devastating epidemics [4, 31, 56] up to the multi-trophic interactions defining ecosystems and microbiomes [50]. Coevolution is defined as the evolutionary change in one antagonist (host) in response to changes in the other antagonist (pathogen). At the genetic level, these changes are determined by genotype x genotype (GxG) interactions between few (up to many) host and pathogen genes. For example, host genotypes differ specifically in their resistance to pathogen strains which in turn differ in their infectivity (ability to infect and cause disease) on the given host genotypes. Host-pathogen GxG interactions are defined by their (i) genetic architecture (how many genes are involved?), (ii) specificity (how many GxG interactions can yield a resistance phenotypic outcome?) and (iii) strength (what is the phenotypic outcome, full resistance up to severe infection?). Knowing the genetic architecture, specificity and strength of GxG interactions is crucial for understanding and predicting the speed and outcome of coevolutionary dynamics [15, 28, 54] and for disease management in agriculture and medicine.

The potentially devastating effects of infection prompted a wealth of genome-wide association (GWAs) studies to identify the genetic architecture (involved genes) of GxG interactions. Single species GWAs are performed by associating genomic variants with a binary disease outcome: (i) infected versus non-infected hosts such as humans [8, 16], invertebrates [11, 12], and plants [22, 44], or (ii) infective/non-infective pathogens [3, 46]. With the growing joint availability of host and pathogen genomic data, two types of joint Genome-Wide Association studies (so-called co-GWAs) have been developed to identify significant GxG loci [9, 10, 40, 58]. Experimental co-GWAs requires a full experimental factorial design of reciprocal infections to assess the outcome of infection (phenotype) [40, 58]. However, controlling for the genetic background and running controlled infection experiments is elusive to human hosts and often difficult to achieve for non-model natural host-pathogen interactions. As an alternative, natural co-GWAs [10, 40] jointly associate genome wide polymorphism data of infected hosts with polymorphism data of their respective infecting pathogen strains [9]. Such natural co-GWAs have since been applied successfully to find associations between human genes and pathogen loci of the Hepatitis C virus (HCV) [5], *Streptococcus pneumoniae* [36] and *Plasmodium falciparum* [7] and to study interactions between *Daphnia magna* host and *Pasteuria ramosa* [23].

Yet, deciphering the specificity and strength of the GxG interactions at the loci of interest has remained empirically out of reach for most host-pathogen systems. Specificity and strength of host-pathogen GxG interactions are classically summarized within the so-called infection matrix, which captures the extent to which each pathogen genotype successfully infects each host genotype (0 meaning full host resistance, and 1 meaning full host susceptibility, Fig. 1a,b). There is a wide range of possible infection matrices which differ in their levels of symmetry, specificity and strength [1, 15, 28]. Further, previous studies suggest that the number of loci involved varies between different host-pathogen systems and there are often epistatic effects between loci [23]. Throughout the article we will focus on four matrices of interest (Fig. 1b): 1) the generalist pathogen (P) infectivity/non-infectivity matrix in which one pathogen genotype has a high infectivity on all host genotypes, 2) the generalist host (H) resistance/susceptibility matrix where one host genotype is resistant against all pathogen genotypes, 3) the specific matching-alleles (MA) matrix where each pathogen genotype is specialized to infect one host genotype as found in the *Daphnia magna - Pasteuria* pathosystem [38], and 4) the specific gene-for-gene (GFG) matrix characterized by an universally infective pathogen genotype and resistance being the result of recognition of a specific pathogen effector allele [55]. GFG interactions are mainly documented for plant-pathogen interactions [25, 55], while MA interactions have been long hypothesized to underlie the interactions between the human major histocompatibility complex (MHC) and most mammalian immunity genes and corresponding pathogen genes [25, 34, 49]. Deciphering the infection matrix via experimental means requires the combinatorial infection assays of many host and pathogen genotypes (clones or isogenic lines with known allelic variants) in controlled conditions, and thus, is prohibitive for most host-pathogen systems (but see [38, 43]).

**Figure 1.**
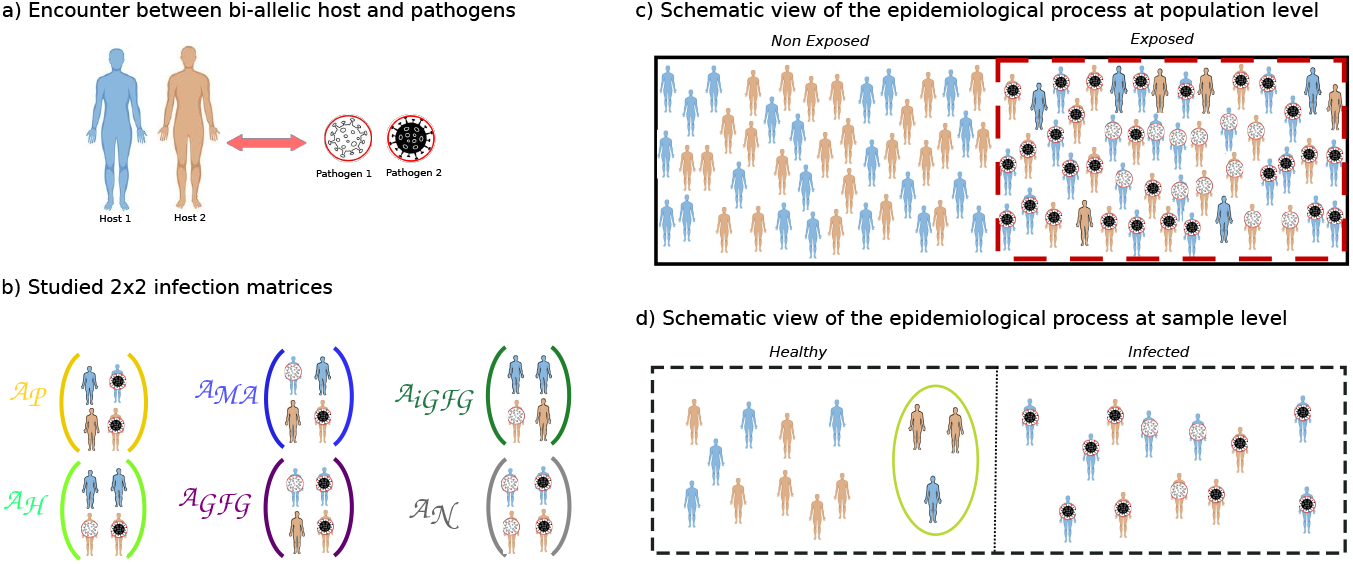
Schematic view of the principles of GxG interactions underlying host-pathogen coevolution and the characteristics of the sampling process. Our model captures the effects of experimental and epidemiological sampling at a single time point of the coevolutionary dynamics. (a) We assume an interaction between a bi-allelic host and a bi-allelic pathogen locus. (b) The outcome of the interaction is summarized by a 2×2 infection matrix with host genotypes as rows and pathogen genotypes as columns. Some classic examples with extreme values are schematically depicted, namely the pathogen infectivity/non-infectivity (𝒜_𝒫_, yellow), matching-allele (𝒜_ℳ𝒜_, blue), inverse gene-for-gene (𝒜_𝒢ℱ𝒢_, dark green), host resistance/susceptibility (𝒜_ℋ_, light green), gene-for-gene (𝒜_𝒢ℱ𝒢_, purple), and neutral (𝒜_𝒩_, grey) matrix. (c) Schematic representation of the infection process and the host’s status at the population level. In a homogeneous population, a proportion *ϕ* (the disease encounter rate) of hosts encounters pathogen infectious propagules at random, and a proportion 1 − *ϕ* does not (epidemiological sampling). Hosts with a solid outline (and circled in green) in the exposed class are resistant to the infection (appear as healthy) due to specific form of the underlying infection matrix. (d) Genomic studies are performed by taking a sample of healthy and infected hosts from the total population, generating a potential bias in sample allele frequencies compared to population alelle frequencies (experimental sampling). On a population level, the frequencies of the host and pathogen alleles are determined by the coevolutionary process (coevolutionary sampling).

We develop here a new theoretical and statistical framework using jointly genomic data of hosts and their pathogens from natural populations to find out the genes underpinning GxG interactions along with inferring the specificity and strength of the interactions. This framework explicitly takes three fundamental sampling processes occurring in any host and pathogen populations into account (Fig. 1c): (i) the (co)evolutionary sampling [25, 39], (ii) the epidemiological sampling, and (iii) the experimental sampling. The first process is a result of coevolution itself, namely that host and pathogen genotype frequencies fluctuate in space and time as a direct result of reciprocal selection (coevolution), genetic drift, mutations and gene flow [28, 54]. As a result, only a subset of all possible interactions between host and pathogen genotypes maybe present at a given point in space and time (Fig. 1c) [25, 54], so that the sampling of host and pathogen genotypes may be incomplete. This effect is a major hindrance for host (or parasite) single species GWAs as it decreases the statistical power when not accounting for the genetic heterogeneity of populations [39]. Second, host genotypes need to encounter corresponding pathogen genotypes in order to get infected as a result of a specific GxG interaction. The frequency of such encounters in natural populations is governed by the allele frequencies, the disease prevalence, the population size and the specific dynamics of host-pathogen encounters. Frankly speaking, an observer cannot know if an uninfected host in a natural population has been exposed to pathogens but is resistant, or if the host has never been in contact with pathogens (Fig. 1c, d). Third, sampling a limited number (subset) of host (infected and non-infected) and pathogen individuals (genotypes) from the entire population for experimental and genomic studies may further blur the true infection matrix (Fig. 1d), because the sampling scheme may over- (or under-) represent some interactions. Altogether, these three sampling processes render the inference of the underlying infection matrix a non-trivial task (Fig. 1). We argue that, so far, the consequences of these stochastic processes on the statistical power of single species GWAs and co-GWAs are poorly understood. Specifically, we expect these three processes to generate variability in the samples’ allele frequencies and in the statistical power of GWAs and co-GWAs studies to detect GxG interactions, depending on the true (yet unknown) infection matrix, epidemiological dynamics, and sampling scheme (infected and/or non-infected hosts) in addition to the previously reported effects of the coevolutionary dynamics and incomplete sampling of hosts and pathogen genotypes [39].

In this study we first derive four different indices based on host-pathogen coevolutionary theory to tackle the problem of inferring the significant GxG interactions and assess their infection matrices under the described stochastic processes. We then incorporate these indices as summary statistics (model based) into an Approximate Bayesian Computation (ABC) framework to analyse jointly genomes of non-infected hosts as well as infected hosts and their matching pathogens. We aim to (i) pinpoint genes underlying GxG interactions, and (ii) infer the underlying infection matrix. As a proof of principle, we infer the interaction matrices underpinning 535 biologically relevant GxG associations between human Single Nucleotide Polymorphism (SNPs) and Singly Amino Acid Polymorphisms (SAAPs) of HCV, using 451 infected individuals and the HCV sequences of the infecting strains [5] complemented by 503 human genomes [2].

## Results

### Indices capture features of the infection matrices

We develop a theoretical model of a temporal snapshot (single-time point) of the outcome of an epidemic process in a host population of large size. In short the model assumes that hosts encounter pathogens at random at a given disease encounter rate *ϕ* (Table S1, Supplementary Text S1). After the infection process the host population can be generally split into two compartments, namely infected hosts (frequency 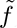) and uninfected hosts (frequency 1 − 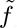). The latter is composed of hosts which either did not encounter pathogens or resisted infection. We denote the frequency of uninfected hosts of type *i* in the entire population as *f_iz_*. Assuming bi-allelic host and pathogen genotypes, there is a maximum of four possible host-pathogen associations in the infected compartment. We denote the frequency of hosts with genotype *i* infected by pathogens genotype *j* in the entire population as *f_ij_* and in the infected subpopulation as 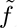*ij*. These frequencies depend on the frequency of hosts of type *i* (*h_i_*), the initial frequencies of pathogen genotype *j* prior to infection, the infection matrix (*α*) and the pathogen encounter rate *ϕ*.

We develop four indices based on co-evolutionary theory to capture the characteristics of a given GxG interaction matrix *α* (Fig. 1b). These indices combine information of host allele frequencies from infected hosts and their associated pathogen strains (pathogen allele frequencies) as in co-GWAs [5, 9, 10] as well as additional information of allele frequencies in a sample of non-infected hosts as in host GWAs [8, 44]. Our first index, the cross-species association (CSA) index, is a cross-species analogue of linkage disequilibrium [27, 40] (also termed interlinkage [23]), which assesses the association between the genotype of infected hosts and the genotype of the respective infecting pathogen strains (thus similar to the natural co-GWAs). The host susceptibility (HS) index compares allele frequencies in the infected versus non-infected host subsamples (thus similar to host GWAs). The pathogen infectivity (PI) assesses the difference between pathogen allele frequencies (thus similar to pathogen GWAs). Finally, the host partitioning (HP) index is designed to reflect the difference of allele frequencies of one host genotype infected by one pathogen allele and when non-infected. The HP index thus contains novel information (compared to co-GWAs and GWAs) on the asymmetry, specificity and strength of the infection matrix.

The four indices are defined as:

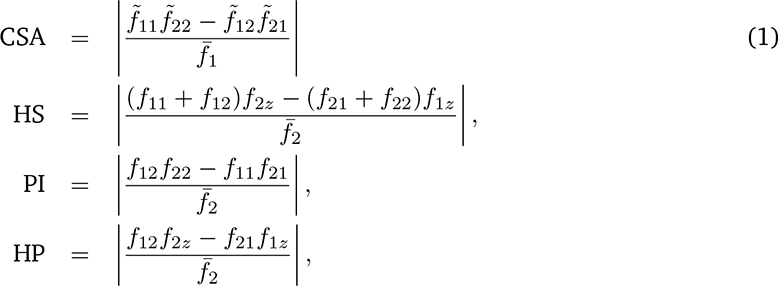

with:

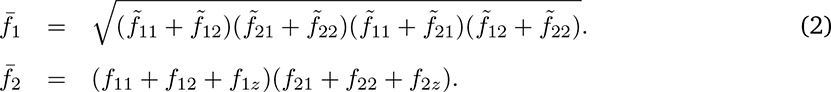

Expressing these indices in terms of the population composition (Table S1) and the coefficients *α_ij_* of an arbitrary 2×2 infection matrix we find (Supplementary Text S1):

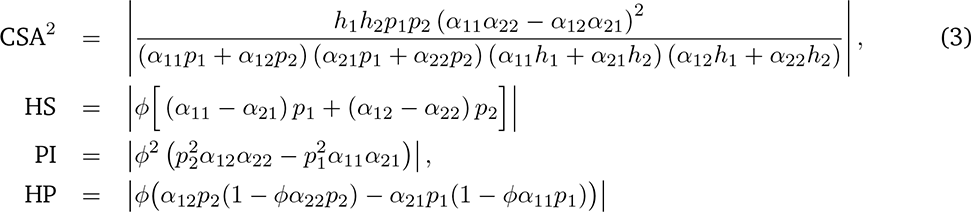

We first derive the population level values of these indices (Table 1, Table S1) for different infection matrices and host and pathogen allele frequencies. This allows us to assess their suitability to distinguish between different matrices. Note that our neutral matrix (all matrix elements 1, Fig. 1b) builds on the hypothesis that the GxG interaction for a given pair of host and pathogen alleles is not relevant for the infection status.

**Table 1.**
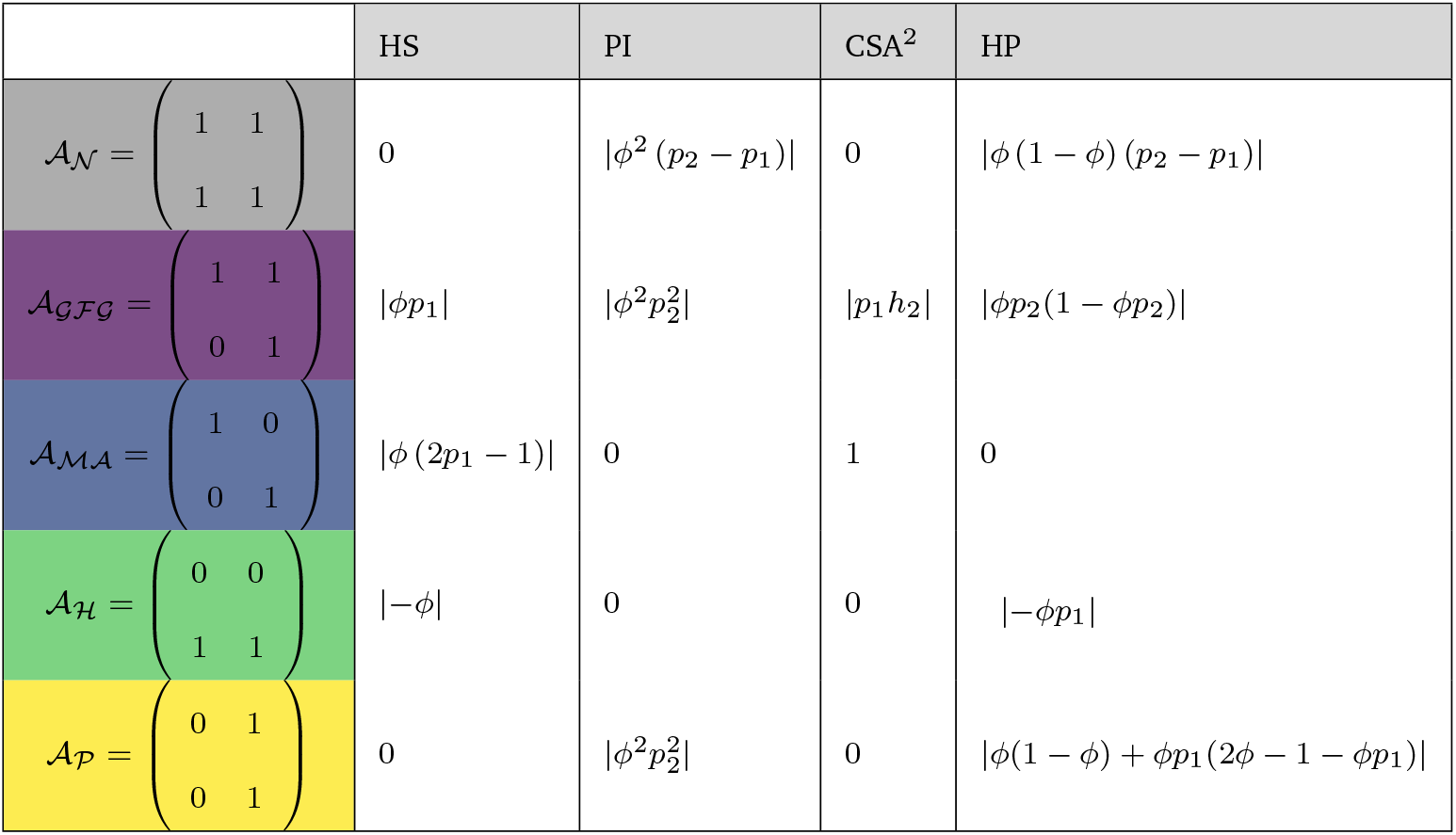
Values of indices for different GxG matrices assuming host genotypes being fully susceptible to infection by pathogen genotype *j* when *α_ij_* = 1 or fully resistant when *α_ij_* = 0.

Studying the most extreme forms (all elements either 0 or 1) of these infection matrices we find that the combination of our indices shows differential behavior among infection matrices. For example the CSA index provides a clear distinction between the GFG and MA matrix from all other matrices. Our results (Table 1) further highlight dependencies of the index values on the disease encounter rate (*ϕ*) and/or non-linear relationships with pathogen allele frequencies prior to host exposure to pathogens (Eq. 3). When we derive expressions of the index values for more general forms of the corresponding GxG matrices (Table S1) the expressions become more cumbersome (Table S2). Yet, we still find that the combination of all four index values shows differential behaviour across the different infection matrices. Therefore, it appears that our four indices, and combinations thereof, can be suitable to discriminate between different types of infections matrices. Extending these theoretical results, it is in principle possible to directly compute the values of the coefficients of the infection matrix (*α_ij_*) by simultaneously solving the set of all equations Eqs. 3. However, this approach shows only reasonable results when the disease encounter rate is known and approximately 50% and when population-level allele frequencies are known (which is in practice not the case because of the effect of the experimental sampling, Fig. 1d, Supplementary Text S1).

### Indices’ behaviour is robust to sampling procedures

We next quantify the behaviour of these four indices to discriminate between different infection matrices under evolutionary, experimental and epidemiological sampling procedures (Fig. 1b, c) (for more details see methods and SI Text). We explore the indices’ distributions over a wide range of minor allele frequencies (*h_i_, p_j_* between 0.05 and 0.5) and allowing for random deviations of the matrix coefficients within a tolerance *δ*. Under evolutionary sampling the ranges of our HP, HS, and PI indices for the entire population are very small for small disease encounter rates, but still distinguishable between different matrices (Fig. 2, top row). The distributions of the indices’ values become more wide spread when taking a sample from the entire population with more or less equal amounts of non-infected and infected individuals (Fig. 2, bottom row). Encouragingly, both for the population and experimental samples, the distribution of indices’ values differs between the different matrices for various disease encounter rates (Fig. 2). More importantly, there is at least one combination of two or more indices (albeit not necessarily linear) for each GxG infection matrix discriminating it from the neutral infection matrix (Fig. 2).

**Figure 2.**
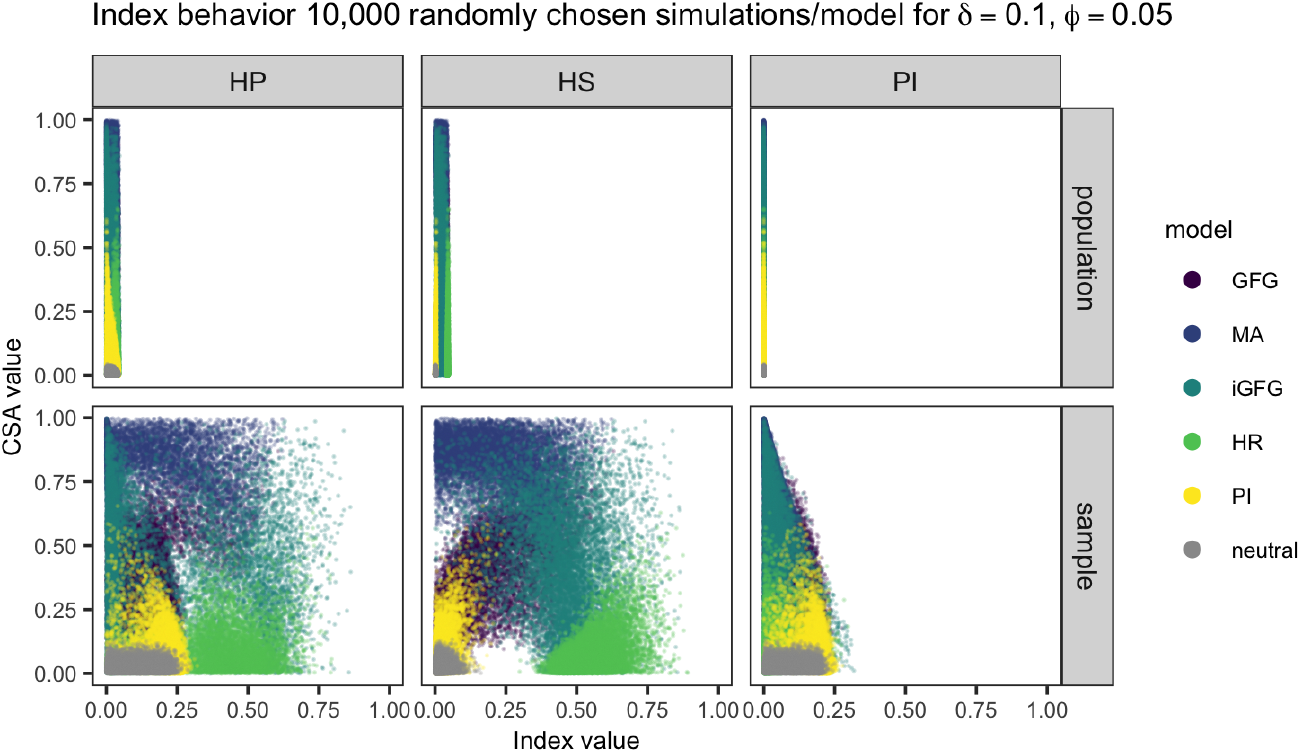
Distribution of values of indices’ pairs comprising CSA (y-axis) and one of the other indices HS, PI or HP (x-axis) for different infection matrices (𝒜_𝒢ℱ𝒢_, 𝒜_ℳ𝒜_, 𝒜_⟩𝒢ℱ𝒢_, 𝒜_ℋ_, 𝒜_𝒫_, 𝒜_𝒩_) for a low disease encounter rate (*ϕ* = 0.05). The population has size *N* = 100, 000 (population row) and a random sample of *n_H_* = 1006 healthy and *n_I_* = 902 infected haploid individuals is taken (sample row). Results are shown for 10,000 simulations where *h*_1_ *∼ U* (0.05, 0.5), *p*_1_ *∼ U* (0.05, 0.5) and *δ* = 0.1. The simulations are a randomly selected subset of the 50,000 simulations used in the ABC model choice.

We observe a strong dependency between the range of possible indices’ values and the disease encounter rate *ϕ* (compare Fig. 2, S1 and S2). Consistent with our theoretical results, the ranges of values for HS, PI and HP are small for low disease encounter rates and increase with higher disease encounter rates (Fig. S1, S2). Yet, for all three disease encounter rates the different matrices are distinguished by the combination of indices under the population sample. As for *ϕ* = 0.05 we observe that taking a fixed sample from the entire population changes the range of observed index values in the sample compared to the population. This effect depends on the specific combination of (host and pathogen) sample sizes and disease encounter rate (Fig. 2, S1-S4). When we consider a sampling scheme with 5% infected and 95% non-infected hosts (keeping the total number of samples to 951 hosts) for a disease encounter rate *ϕ* = 0.05, the distribution of indices’ values becomes more narrow and more similar to that of the population sample. This potentially decreases the extent to which different matrices can be discriminated (Fig. S4). On the other hand if we consider a sampling scheme with 95% infected and 5% non-infected hosts (keeping the total number of samples to 951 hosts) the range of indices’ values further broadens and becomes less similar to the population sample. The difference in sample indices’ distributions reflects the effect of experimental sampling of infected hosts on top of the epidemiological sampling for a low disease encounter rate. Our results exemplify the, so far, largely ignored effects of the epidemiological and experimental sampling in natural co-GWAs and the importance of deriving of optimal sampling schemes to overcome this interplay.

Increasing the tolerance threshold (value of *δ*) increases the amount of overlap between the indices’ distributions. As a consequence different matrices may be confounded (Fig. S5 for *δ* varying between 0.1 and 0.3). In other words, choosing a low tolerance parameter generates a more stringent statistical test to disentangle between the neutral infection matrix and other matrices and should decrease the rate of false positives (association and underlying matrices appearing to be biologically relevant whereas these are in fact neutral).

### An ABC framework allows to infer the infection matrices

We then use these simulation results (*ϕ* = 0.05 and *δ* = 0.1) in an Approximate Bayesian Computation (ABC) framework to infer the infection (GxG) matrix (neutral, MA, GFG,…) for a given association. We use our four indices as ABC summary statistics. A GxG association is not biologically relevant (significant) for a host-pathogen interaction if the ABC model choice procedure reveals the neutral matrix as the best (or equally best) model. We assess the statistical power of ABC model (matrix) choice by running a leave-one-out cross-validation (rejection algorithm, tolerance=5%) based on randomly chosen 500 simulations per infection matrix and inferring the best model using simulations for all matrices (50,000 per matrix). We demonstrate that our ABC based on our four indices can discriminate between all matrices (Table 2, S3, S4), especially between biologically relevant GxG matrices and the neutral matrix (under the most stringent threshold, Table S3, S4). The host resistance and the MA matrix can be well discriminated from all other matrices, whereas the pathogen infectivity and iGFG matrices may still be confounded with other matrices (Table 2). Therefore, accounting for sampling effects (as introduced in priors for various parameters and fixed sample sizes) within our ABC framework allows to disentangle between the different infection matrix models (Table 2, S3, S4).

**Table 2.** Results of a leave-one-out ABC (rejection) cross-validation for 500 randomly chosen simulations per infection matrix under low disease encounter rate. For each model 50,000 simulations are produced for *h*_1_ *∼ U* (0.05, 0.5), *p*_1_ *∼ U* (0.05, 0.5), *δ* = 0.1, *ϕ* = 0.05, *N* = 100, 000, *n*_I_ = 902 haploid and *n_H_* = 1006 haploid.

| True model | Inferred model |  |  |  |  |  |
| --- | --- | --- | --- | --- | --- | --- |
|  | neutral | GFG | MA | iGFG | HR | PI |
| neutral | 458 | 0 | 0 | 0 | 0 | 42 |
| GFG | 16 | 361 | 24 | 46 | 6 | 47 |
| MA | 0 | 7 | 471 | 22 | 0 | 0 |
| iGFG | 2 | 52 | 63 | 244 | 138 | 1 |
| HR | 0 | 0 | 0 | 7 | 492 | 1 |
| PI | 114 | 39 | 0 | 0 | 0 | 347 |

### 535 biologically relevant GxG associations between humans and HCV

We now apply our ABC framework combining a dataset of human diploid host sequences and their infecting HCV (hepatitis C virus) strains [5] (the infected sample) and additional sequences from the 1,000 genomes project [2, 52] (the non-infected sample). In order to limit confounding effects of population structure in the data, we restrict our analysis to the subset of 451 individuals of European ancestry (PCA in Fig. S6, [5]). As previously described [5], we convert the viral nucleotide sequence data into Single Amino Acid Polymorphisms (bi-allelic SAAPs) data. Our non-infected sample consists of 503 diploid individuals of European ancestry from the 1,000 genomes project (filtering SNPs corresponding to those from [5], PCA in Fig. S6). We filter for a minor allele frequency (MAF) MAF *>* 0.2 to maximize the power to disentangle between infection matrices. As highlighted above, below this frequency, several stochastic sampling effects decrease significantly the power to pinpoint relevant GxG associations. We compute our four indices for all possible pairwise associations between 326,520 human SNPs and 208 SAAPs. For the 800 top associations defined as exhibiting the highest values of our indices, we run the ABC model choice between the possible six infection matrices (neutral, GFG, iGFG, MA, H, P). Our model choice results in 535 interactions which differ from the neutral matrix based on a Bayes factor threshold of two (BF *>* 2) and for a matrix tolerance threshold *δ* = 0.1 (Fig. 3). For each of the 535 associations, we infer the most probable infection matrix (Table S5-S8).

**Figure 3.**
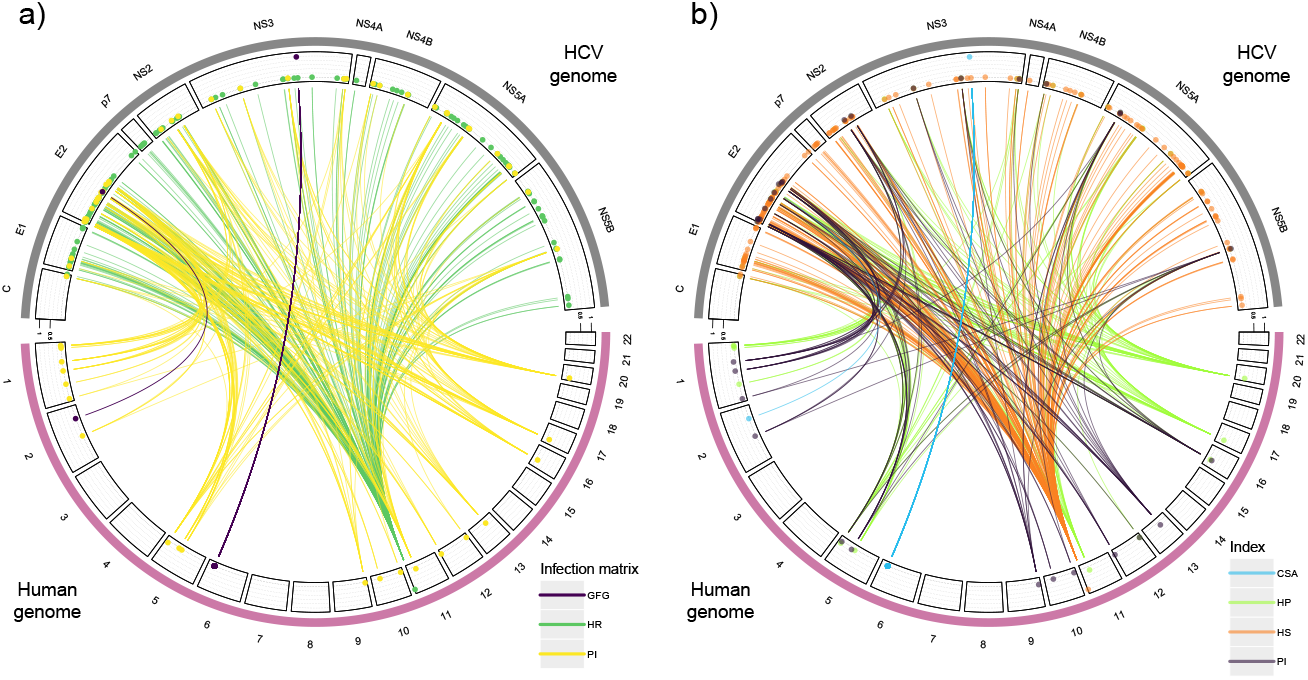
Genome-to-genome relevant associations from the ABC-model choice for all 535 associations (*BF >* 2 to the neutral matrix). a) The 535 associations between host SNPs and pathogen SAAPs colored by the single best infection matrix. b) The 535 associations colored by the most informative index. The human chromosomes are shown on the bottom, the virus contigs on the top. The second circle of lines indicate the number of sites sharing one association (most inner line indicates a single site up to most outer line indicating multiple closely linked sites). The color coding is as follows for the infection matrices: purple=𝒜_𝒢ℱ𝒢_, darkgreen=𝒜_ℋ_, yellow=𝒜_𝒫_. For the indices blue is for CSA-index, lightgreen for HP-index, orange for HS-index, and purple for PI-index.

We summarize the estimated infection matrices and their distributions by index (Fig. 3, S7, S8). We find two main groups of associations with an estimated gene-for-gene matrix (𝒜_𝒢ℱ𝒢_): one group includes several SNPs on the human chromosome 6 falling into the MHC region and the HCV gene NS3, and the second group has only one association under GFG between one SNP at the clathrin heavy chain linker domain containing 1 (CLHC1) gene on chromosome 2 and an SAAP on the HCV gene E2. Furthermore, we find several associations with an estimated resistance matrix (𝒜_ℋ_) between one position at the Lymphocyte-specific protein 1 (LSP1) on chromosome 11 and SAAPs at various viral genes. Finally, we find also several pathogen infectivity matrices (𝒜_𝒫_) between 21 SNPs in the human genome and 45 SAAPs in the HCV genome. We also highlight, that we do not find any associations which are indicative of a matching-allele (MA) infection matrix, even when lowering the detection threshold (higher *δ*) and considering competing best models (Tables S5-S8). Analyzing the details of these 535 biologically relevant GxG associations, we find few host sites (106) especially exhibiting GFG or resistance matrices, while pathogen AA (221) exhibit chiefly infectivity matrices. We also compare our results to a co-GWAs on the subset of 451 European human infected individuals (and their 451 pathogen strains) following the previous analysis [5] using plink. Our CSA index shares all associations with our Bonferroni corrected co-GWAs (Fig. S9). In addition, 68 candidates from our European CSA index also appear in the 104 top candidates from the Bonferroni corrected results obtained previously for the full data set (Fig. S10, [5]). We conclude that our ABC framework based on our CSA index reveals relevant GxG associations but is more stringent than co-GWAs studies. Using four indices which capture different aspects of infection matrices, we are able to reveal new associations which were not previously reported, especially potential human resistance alleles to HCV (under GFG and resistance matrices) and HCV infectivity alleles (under infectivity matrix).

## Discussion

We derived four indices to tackle the problem of inferring the underlying infection matrix from host-pathogen association data. Further, we developed some general predictions on the behavior of these indices, establish their joint ability to discriminate between several infection matrices and use them successfully as summary statistics in an ABC framework to reveal infection matrices in a human host/HCV virus data set. Therefore, our study is not only the first study attempting to establish the theoretical aspects of natural co-GWAS and the effect of various type of sampling, but also lays ground for a framework to infer the infection matrix using natural co-GWAs set-ups. We specifically present results tuned for studying the interaction between European humans and HCV, namely we assume a sample size of 451 infected diploid humans (and their respective viral strains) and 503 non-infected European humans, and a known disease encounter rate of 0.05 (slightly higher than the disease prevalence of approximately 3% previously reported [42, 45]).

Our Bayesian framework identifies genes that exhibit statistically relevant associations which have some support from previous studies. All observed associations at the major histocompatibility complex (MHC) on chromosome 6 are associated with one viral site located at position 1,444 on the non-structural protein 3 (NS3) gene. All these human sites are likely linked to alleles at the HLA genes due to the high amount of linkage disequilibrium across the region [6]. The HLA genes of a patient determine which viral peptides are presented to T cells as part of the adaptive immune response. This process can drive viral evolution and result in the emergence of viral escape mutations, which have been previously identified for the NS3 gene [5, 41]. There is some empirical evidence that small interfering RNA-mediated clathrin heavy chain depletion affects endocytosis of HCV [14, 20]. Therefore, we speculate that the CLHC1 (Clathrin Heavy Chain Linker Domain Containing 1) gene may be involved in such process, possibly supporting our finding of a GFG interaction of this gene with an amino acid in the HCV gene E2. We further found a putative resistance allele at the Lymphocyte-specific protein 1 (LSP1) gene which is an F-actin binding protein. This protein is involved in the regulation of various immune system functions, including lymphocyte activation, proliferation, and migration [47]. It also has been shown to play a role in endocytosis and transendothelial migration of leukocytes, allowing these to be recruited to the sites of inflammation [37, 57]. Studies demonstrating that the depletion of LSP1 significantly reduces the rate of endocytosis of HIV particles [18, 19], could suggest that this protein may also play a role in the endocytosis of HCV.

Despite likely coevolving with humans over thousands of years in Africa, HCV has a very recent history of infection and spreading in the European human population (Fig. S11, [24, 26]). Therefore, the biologically most relevant SNP-SAAP associations should be interpreted in the light of the HCV virus adapting to existing standing genetic variation in the European population within the (approximately) last 150 years. Experimental results from a bacteria-phage coevolution interaction [33], indicate that initial coevolutionary dynamics are characterized by rapid fixation of advantageous alleles in hosts and pathogens (arms race dynamics [13]). The dynamics are then replaced by trench warfare dynamics [51] with maintenance of two or more alleles at the coevolving genes by balancing selection. Our inferred asymmetric matrices (host resistance, pathogen infectivity and GFG) likely indicate that we capture the initial dynamics of the interaction between humans and HCV in Europe. Asymmetric matrices are more likely to generate arms race dynamics especially when population sizes are small [1, 53, 54]. In this light, we interpret the finding of a resistance matrix at the LSP1 gene as an indication that resistance to HCV may be segregating in the human population. Several mutations in the virus populations have likely been selected for overcoming this resistance allele (green lines in Fig. 3). In addition, several SAAPs with inferred infectivity matrices likely indicate that strains of HCV exhibit mutations allowing them to infect and match several host genes and alleles. In other words, there are virus strains with different infectivity ranges. Finally, the inferred GFG interactions indicate that the virus has evolved to overcome host recognition alleles at several MHC genes and at one gene on chromosome 2 (CLHC1). These human alleles likely provided initial resistance to HCV at the onset of the epidemics which has been overcome by subsequent mutations in the virus.

We speculate that further extending our inference framework to data sampled from different time points can potentially help to elucidate the speed and timing of coevolution and changes in the GxG interactions at the genetic level. One key prediction from co-evolutionary theory is that due to various stochastic and selective processes, the number of genes under coevolution and the corresponding infection matrices are subject to change over time [15, 25], exhibiting various degrees of asymmetry [1, 28]. This in turn generates different coevolutionary dynamics in time (arms race and trench warfare dynamics, respectively, [29, 54]). It is possible that in the long term run coevolution between HCV and other human populations, SNPxSAAP interactions may exhibit different underlying infection matrices promoting the occurrence of trench warfare dynamics and balancing selection, namely : (i) symmetric MA interactions, or (ii) asymmetric GFG interactions with the necessary, but not sufficient, [53, 54] condition that costs of resistance and infectivity exist at these coevolving loci. Applying our inference framework to other diseases with a range of short to long term coevolutionary histories would shed light on the speed of coevolution between humans and their viruses [30] and the underlying coevolutionary dynamics.

We specifically focus on random disease transmission and low disease encounter which likely best describe HCV transmission dynamics in Europe [42, 45]. This allows us to account for the effect of the epidemiological sampling on the distribution of allele frequencies in the population and in our experimental samples, without the need to specify a corresponding prior for the disease encounter rate in the ABC. Our use of priors for host and pathogen allele frequencies takes into account that the ’true’ allele frequencies prior to the infection process (*h_i_* and *p_j_* in our model) are unknown. The observed allele frequencies in the sample represent indeed the outcome of the joint interaction of disease encounter rate, the ’true’ but hidden allele frequencies and the infection matrix. We acknowledge that disease encounter rates might be less well known for host-pathogen interactions involving non-model species. One way to tackle this limitation for non-model plant host-pathogen interactions would be to obtain estimates for the range of disease encounter rates from field data and include this range as an additional prior into the ABC simulations. However, based on our analytical results, we expect this approach to be only successful if the corresponding estimated range of the disease encounter rate is relatively narrow.

Furthermore, we follow previous GWAs and co-GWAs approaches and assume sufficiently large sample sizes (several hundred individuals) to allow the detection of significant associations. As some of our indices rely on estimating the allele frequencies in the non-infected sub-sample and the infected sub-sample with comparatively small error, it is crucial to obtain a sample that well reflects the population frequencies of genotypes/phenotypes in the entire population. Specifically, if the disease encounter rate is small, it is important to sample well enough the infected part of the population. Conversely, if the disease encounter rate is high, sufficient sampling of the non-infected part of the population becomes important. This emphasizes the importance of taking care the interaction between sampling size and disease prevalence into account when devising sampling schemes in co-GWAS studies. Especially low sample sizes, are very likely to produce biased allele and association frequencies in the sample and hence, erroneous infection matrix estimates.

An inherent difficulty for any co-GWAs and our ABC is to confidentially detect associations which involve alleles with low frequencies. Therefore, we conservatively restricted our testing to loci with a minor allele frequency (MAF) *>* 0.2 to avoid an excess of false positives. However, coevolutionary dynamics can transiently decrease allele frequencies or maintain alleles at low frequencies as a result of negative indirect frequency-dependent selection [39, 53, 54]. Therefore, we speculate that our ABC method can be further improved by incorporating sample allele frequencies as additional summary statistics and by using association data from several time points. We expect the later to help with better tracking the allele frequency changes over time which directly result from the coevolutionary dynamics and the underlying infection matrix. One further current limitation of our model (and all co-GWAs) is not accounting for epistatic effects and multi-locus infection matrices. Indeed, in some species such as *Daphnia*, the resistance phenotype depends on epistatic interactions between several loci [38]. By integrating such knowledge of epistatic interactions, the results of the co-GWAs could recently be improved and additional genes of interactions discovered [23]. Integrating time-sampled data, more summary statistics and the effect of epistasis between host (or pathogen) loci into our framework constitute the topic of future work.

It is well known from the GWAs literature that spatial structure in the host and pathogen samples can affect and distort the power to detect associations. Therefore, we restricted our analysis to a single population (European) without any obvious population structure. Our results align with, and are more conservative, than the previous co-GWAs [5] applied to the same data set (Fig. S9-S10). Therefore, we are confident that our framework is stringent and exhibits a low rate of false positives. In addition, recent studies demonstrate the usefulness of using local population rather than widespread sampling in GWAS setting [32]. Accounting for spatial structure covariates and kinship matrix is another topic of future work.

In conclusion, we built here an ABC integrative method based on four indices as summary statistics which combines ideas from host or pathogen GWAs as well with host-pathogen co-GWAs and additional information from non-infected hosts. Our framework is based on a widely applicable theoretical infection model, takes into account various sampling procedures defining observed host and pathogen allele frequencies, and thus allows us to define a threshold to detect biologically relevant GxG associations in a Bayesian framework. Our model and framework should be also applicable to other GxG interactions, such as between hosts and mutualistic symbionts or between chloroplasts/mitochondria x nuclear genes interactions.

## Methods

### Definition of indices

The CSA index is calculated based on the frequencies of host/pathogen genotype combinations in the infected subpopulation/sample. We define the frequency of host genotype *i* infected by pathogen genotype *j* among all infected individuals as 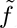*ij* (*i, j ∈* [1, 2]).

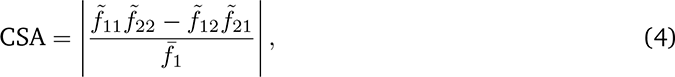

By analogy with the linkage disequilibrium measure in population genetics, we normalize the index by the square root of the product of all infected host and pathogen allele frequencies.

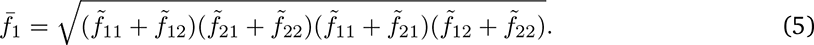

We define the genotype frequencies of uninfected hosts of type *i* in the population/sample as *f_iz_*. Individuals can be uninfected due to two reasons: (i) they have not been exposed to the pathogen *f_i_*_0_, or (ii) they had a pathogen encounter but resisted infection *f_i_*_3_. We lump these two frequencies into a single frequency *f_iz_* as in a natural population it is usually impossible to tell apart the difference.

The HS, PI and HP indices are defined as follows:

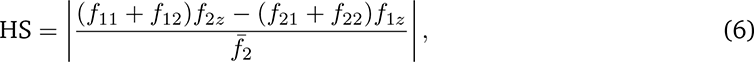

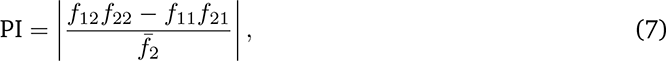

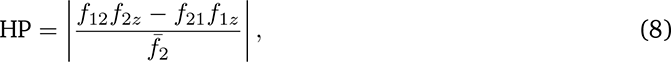

With

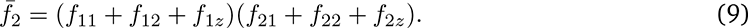

We derived expressions for these indices for a single point in time given initial host genotype frequencies *h_i_*, pathogen genotype frequencies *p_j_*, a disease encounter rate *ϕ* and a given infection matrix *α* (see Supplementary Text S1).

### Stochastic simulations

Next we assessed the effect of three types of stochastic processes on the behavior of the indices for all matrices in Table 1: (i) deviations of the matrix elements from the extreme values 0 and 1 and varying host and pathogen alleles frequencies, (ii) random sampling of a fixed number *n_H_* healthy and *n_I_* infected individuals from a population of size *N*, and (iii) a small (*ϕ* = 0.05), intermediate (*ϕ* = 0.5) or large disease encounter rate (*ϕ* = 0.95). Therefore, we developed a simple host-pathogen interaction model/simulator for a population of size *N* = 100, 000, which is subdivided into a compartment of hosts interacting with the pathogen and a compartment not interacting with the pathogen based on a disease encounter rate *ϕ*. Hosts encounter pathogens in a frequency dependent manner and upon encounter between a host of type *i* and a pathogen of type *j* the host gets infected with probability *α_ij_*. We run simulations for the six infection matrices in Table 1. Therefore, we first randomly chose one of the possible assignments of *α* values (0 or 1) to the matrix elements *α_ij_* for the given matrix (two possibilities for MA, HR, PI and four possibilities for GFG, iGFG, see Supplementary Text S1). Second, after assigning 0s or 1s to the matrix elements, we replaced each element *α_ij_* = 1 by randomly drawing a value from a corresponding uniform distribution *U*_[1_*_−δ,_*_1]_, and we replaced each element *α_ij_* = 0 by drawing from a uniform distribution *U*_[0_*_,δ_*_]_ (Supplementary Text S1). The initial host frequencies *h*_1_ and pathogen *p*_1_ for each simulation are both drawn from a uniform distribution *U* (0.05, 0.5). Based on the resulting matrix and initial host and pathogen frequencies, we calculated the frequencies of all possible infected *f_ij_* and healthy *f_iz_* host phenotypes in the entire population and the respective sub-population (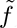*ij* for infected, 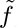*iz* for healthy) (equations in Table S2). We then randomly picked a sample of *n_I_* = 902 haploid infected individuals (drawn from a multinomial distribution *Mult*(*n_I_*, 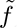_11_, 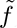_12_, 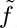_21_, 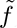_22_)) and *n_H_* = 1006 hap-loid healthy individuals (drawn from a binomial distribution *B*(*n_H_*, 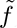_1*z*_)). Following this approach we generated 50,000 simulations for each matrix for each combination of *ϕ* ∈ {0.05, 0.5, 0.95} and *δ* ∈ {0.1, 0.2, 0.3}.

### ABC Leave-one-out cross validation for model selection

We first run a leave-one-out cross-validation using our simulated data set to test the suitability of ABC model choice with our four indices as summary statistics to distinguish between the six different matrices. Leave-one-out cross-validation was run separately for each combination of *ϕ* and *δ* for a cross-validation sample of size 500 using the function `cv4postpr` in the R-package abc (rejection algorithm, tolerance=0.05) [21].

### Application to human data

In the next step we combined two existing human data sets to apply and test our framework. For the infected sample, we used human genome-wide genotype data and HCV whole-genome sequence data from [5]. This data was collected from a total of 541 patients infected by HCV genotypes 2 and 3. We only used a subset of 451 humans of European ancestry to prevent confounding effects of population structure. For the pathogen genome information, we used the viral (nucleotide and protein) data from [5] from NCBI GenBank (accessions KY620313–KY620880). Following [5], we generated whole-genome viral consensus sequences (nucleotide and protein) for each patient using MAFFT (v.7.429) [35]. Future details of how we processed the virus data for our analysis are given in Supplementary Text S1. For the non-infected sample, we used genotype data from the 1,000 Genomes Project Phase 3 [2]. We used the 503 samples from five sub-populations of European ancestry and therefore, retrieved vcf-data from 91 individuals from England and Scotland (GBR), 99 Finnish individuals (FIN), 99 Utah residents with Northern and Western European ancestry (CEU), 107 Spanish individuals (IBS) and 107 Italian individuals (TSI) for a total of 503 genomes (details in Supplementary Text S1).

### Co-GWAS

We run a natural co-GWAS with PLINK2 ([48], [17]) on the data using a logistic-regression with the firth-fallback option. For each regression, we used the presence of a particular amino acid at a given position in the viral alignment as a response variable and the genotype at a given human SNP as the genotype. To account for multiple testing we calculated several p-value adjustments using the −−*adjust* option of PLINK2. We incorporate Sex, human PC1-PC3 and virus PC1-PC10 as covariates in the PLINK co-GWAs.

### Index calculation with application to the HCV data

We obtained frequencies for each host-virus association from the infected human data set using PLINK2 and vcftools. We also extracted the frequencies of alleles in the non-infected human sub-sample. Combining these frequencies, we calculated all of our four indices using the equations 4,6,7,8 with customized R-scripts. After that, we retrieved a summary table with the top outlier associations for each index.

### Model choice for the top association candidates

The authors thank Joy Bergelson, Dieter Ebert and Lars Raberg for comments on an earlier version of the manuscript. AT acknowledges support from the Deutsche Forschungsgemeinschaft project numbers 274542535 (TE809/3) and 221691301 (TE809/6) within the Priority Program SPP1819 ”Rapid adaptation”. The data aquisition was funded by a grant from the Medical Research Council (MRC) (MR/K01532X/1; to the STOP-HCV Consortium). E.B. is funded by the NIHR Oxford BRC. M.A.A. is supported by a Sir Henry Dale Fellowship jointly funded by the Royal Society and the Wellcome Trust (220171/Z/20/Z).

### Data availability

All codes and pipelines developed for index computation and ABC framework, are available at https://gitlab.lrz.de/population genetics/cogenomics method. The genomic data can be obtained upon request to the STOP-HCV consortium (https://www.expmedndm.ox.ac.uk/stop-hcv).

## Supporting information

Sifigures-SItables

SItext-methods

## Author contributions

HM, SJ and AT designed the project; HM, SJ and AT developed the model; HM, SJ and LM performed the simulations, LM performed the data analysis, HM, SJ and LM wrote the manuscript with support from AT. STOP-HCV consortium, MAA and VP obtained and pre-processed the sequence data before making them available. The authors declare no conflict of interest.

## STOP-HCV Consortium: list of members and affiliations

Eleanor Barnes, Emma Hudson, Paul Klenerman, Peter Simmonds (Nuffield Department of Medicine and the NIHR Oxford BRC, Peter Medawar Building for Pathogen Research, University of Oxford, Oxford, UK); Chris Holmes (Department of Statistics, University of Oxford, Oxford, UK); Graham Cooke (Wright–Fleming Institute, Imperial College London, London, UK); Geoffrey Dusheiko (Institute of Liver Studies, King’s College Hospital NHS Foundation Trust, London, UK); John McLauchlan (MRC–University of Glasgow Centre for Virus Research, Glasgow, UK); Mark Harris (School of Molecular and Cellular Biology, Faculty of Biological Sciences and Astbury Centre for Structural Molecular Biology, University of Leeds, Leeds, UK); William Irving (University of Nottingham, Queen’s Medical Centre, Nottingham, UK); Philip Troke (Gilead Sciences Ltd., London, UK); Diana Brainard and John McHutchinson (Gilead Sciences, Foster City, CA, USA); Charles Gore and Rachel Halford (Hepatitis C Trust, London, UK); Graham R Foster (Queen Mary University of London, London, UK); Cham Herath (Gilead Sciences, Middlesex, UK).

## Notes

### Competing Interest Statement

The authors have declared no competing interest.

