## Supplementary material for "Inference of host-pathogen interaction matrices from genome-wide polymorphism data": Sifigures-SItables

### Supplementary Figures

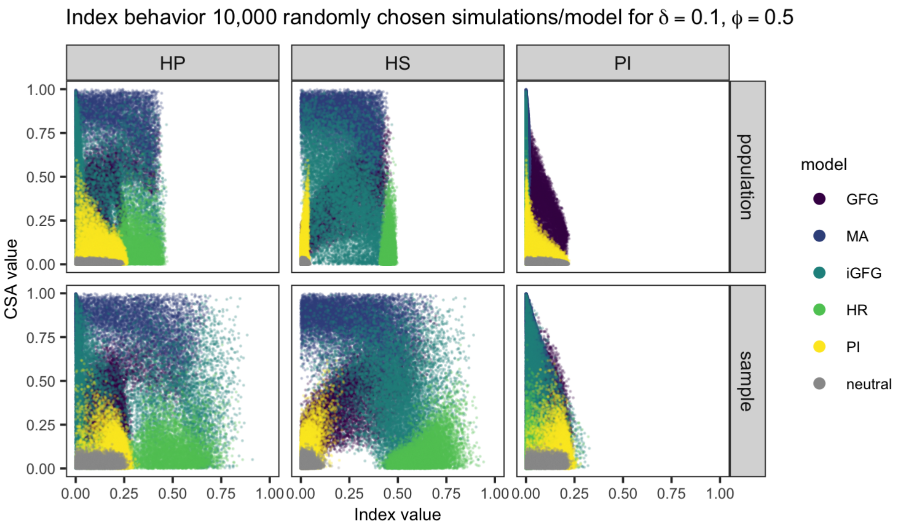


Figure S1  Effect of an intermediate disease encounter rate of $\phi = 0.5$ on the observed index values for different infection matrices ($\mathcal{A}_{\mathcal{GFG}}$, $\mathcal{A}_{\mathcal{MA}}$, $\mathcal{A}_{i\mathcal{GFG}}$, $\mathcal{A}_{\mathcal{H}}$, $\mathcal{A}_{\mathcal{P}}$, $\mathcal{A}_{\mathcal{N}}$). The population has size $N = 100,000$ and a random sample of $n_H = 1006$ healthy and $n_I = 902$ infected haploid individuals is taken. Results are shown for 10,000 simulations where $h_1 \sim \mathcal{U}(0.05, 0.5)$, $p_1 \sim \mathcal{U}(0.05, 0.5)$, $\delta = 0.1$ and $\phi = 0.5$.

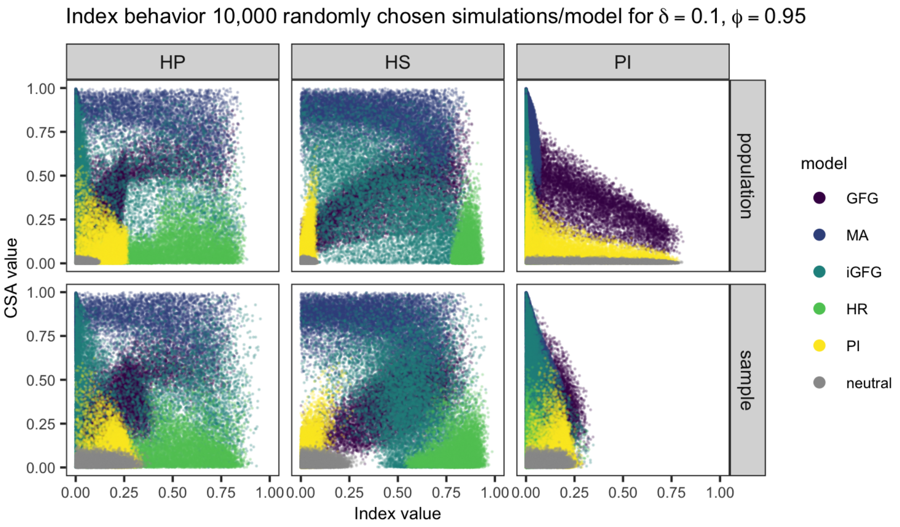


Figure S2  Effect of a high disease encounter rate of $\phi = 0.95$ on the observed index values for different infection matrices ($\mathcal{A}_{\mathcal{GFG}}$, $\mathcal{A}_{\mathcal{MA}}$, $\mathcal{A}_{i\mathcal{GFG}}$, $\mathcal{A}_{\mathcal{H}}$, $\mathcal{A}_{\mathcal{P}}$, $\mathcal{A}_{\mathcal{N}}$). The population has size $N = 100,000$ and a random sample of $n_H = 1006$ healthy and $n_I = 902$ infected haploid individuals is taken. Results are shown for 10,000 simulations where $h_1 \sim \mathcal{U}(0.05, 0.5)$, $p_1 \sim \mathcal{U}(0.05, 0.5)$, $\delta = 0.1$ and $\phi = 0.5$.

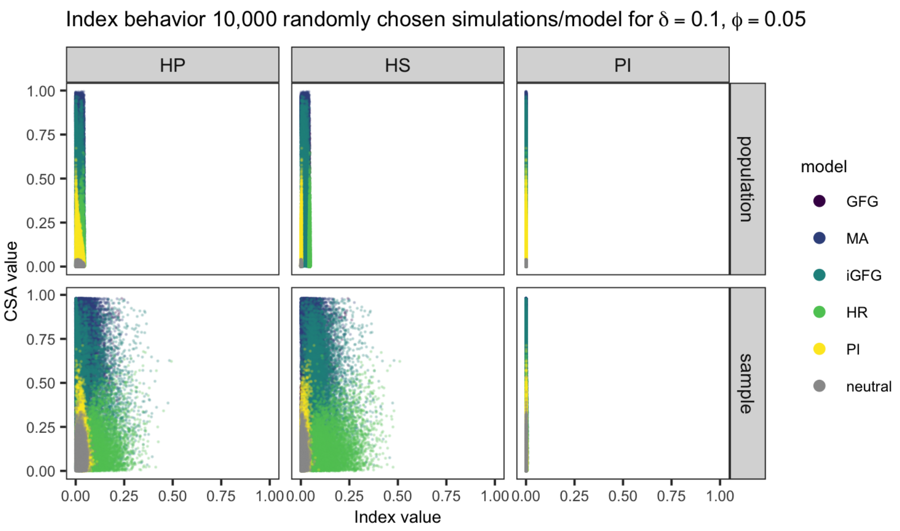


Figure S3 Effect of different sample sizes of $n_H = 1813$ healthy and $n_I = 95$ infected haploid individuals on the observed summary statistics for different infection matrices ($\mathcal{A}_{\mathcal{GFG}}$, $\mathcal{A}_{\mathcal{MA}}$, $\mathcal{A}_{i\mathcal{GFG}}$, $\mathcal{A}_{\mathcal{H}}$, $\mathcal{A}_{\mathcal{P}}$, $\mathcal{A}_{\mathcal{N}}$) in a population of size $N = 100,000$. Results are shown for 10,000 simulations where $h_1 \sim \mathcal{U}(0.05, 0.5)$, $p_1 \sim \mathcal{U}(0.05, 0.5)$, $\delta = 0.1$ and $\phi = 0.05$.

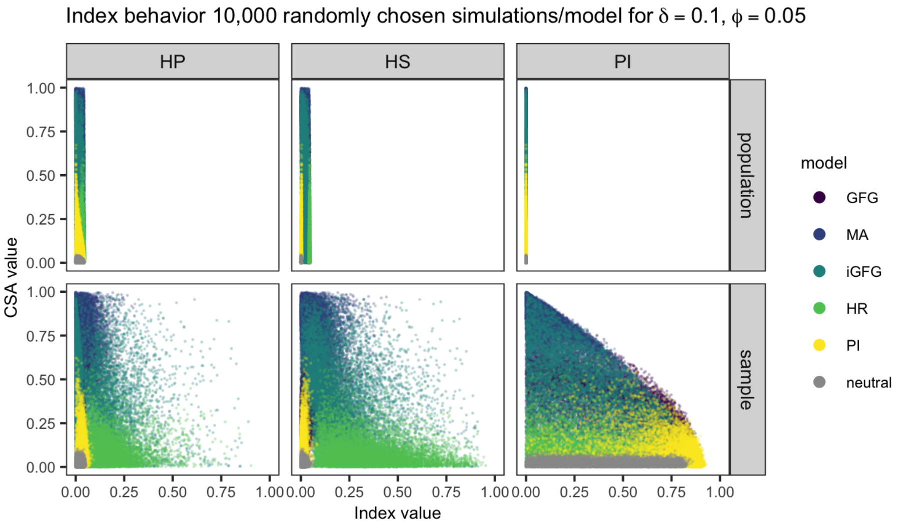


Figure S4 Effect of different sample sizes of $n_H = 95$ healthy and $n_I = 1813$ infected haploid individuals on the observed summary statistics for different infection matrices ($\mathcal{A}_{\mathcal{GFG}}$, $\mathcal{A}_{\mathcal{MA}}$, $\mathcal{A}_{i\mathcal{GFG}}$, $\mathcal{A}_{\mathcal{H}}$, $\mathcal{A}_{\mathcal{P}}$, $\mathcal{A}_{\mathcal{N}}$) in a population of size $N = 100,000$. Results are shown for 10,000 simulations where $h_1 \sim \mathcal{U}(0.05, 0.5)$, $p_1 \sim \mathcal{U}(0.05, 0.5)$, $\delta = 0.1$ and $\phi = 0.05$.

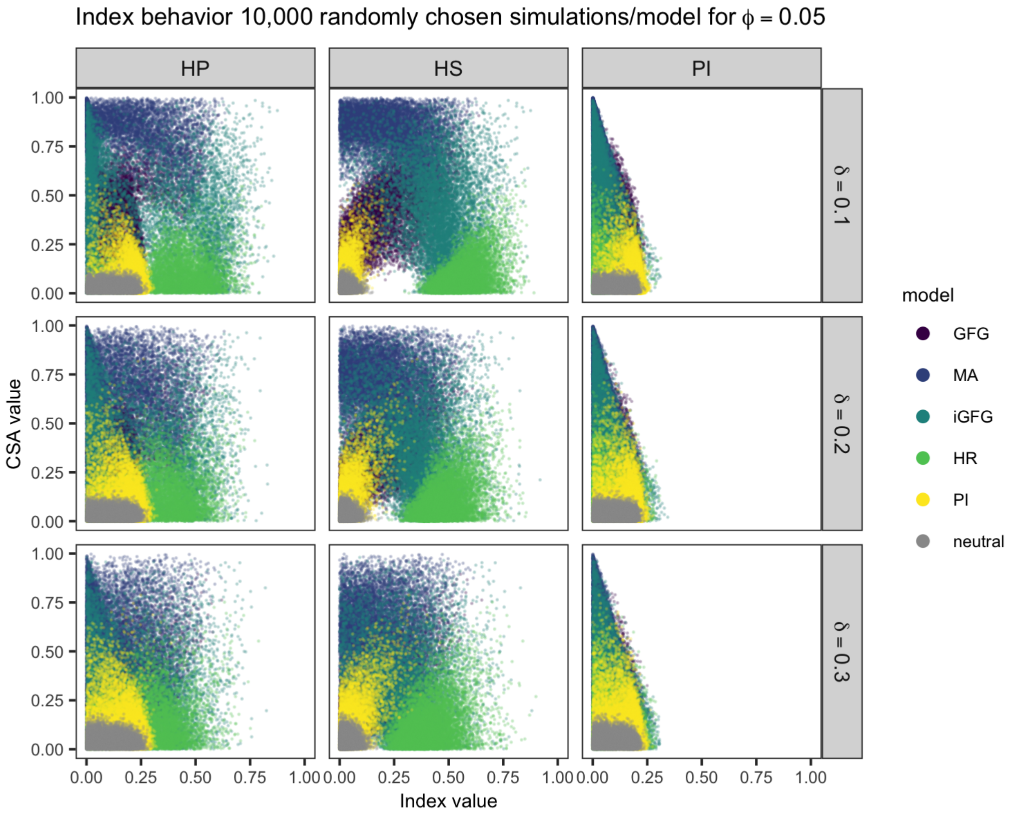


Figure S5 Effect of different values of the parameter $\delta$ (0.1, 0.2 and 0.3) affecting the matrix coefficients of a given matrix on the observed summary statistics for different infection matrices ($\mathcal{A}_{\mathcal{GFG}}$, $\mathcal{A}_{\mathcal{MA}}$, $\mathcal{A}_{i\mathcal{GFG}}$, $\mathcal{A}_{\mathcal{H}}$, $\mathcal{A}_{\mathcal{P}}$, $\mathcal{A}_{\mathcal{N}}$) in a population of size $N = 100,000$. Results are shown for 10,000 simulations where $h_1 \sim \mathcal{U}(0.05, 0.5)$, $p_1 \sim \mathcal{U}(0.05, 0.5)$ and $\phi = 0.05$.

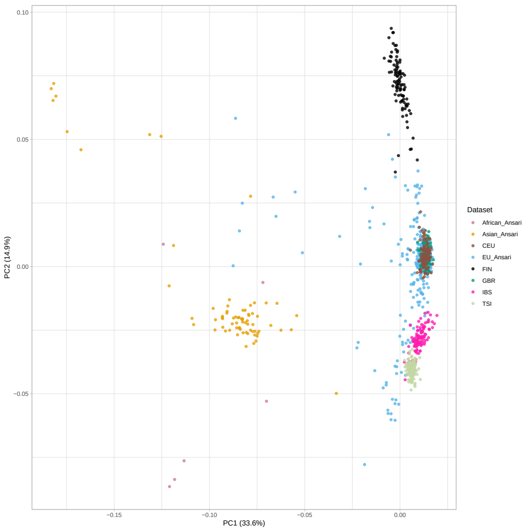


(a) PCA human data for all populations

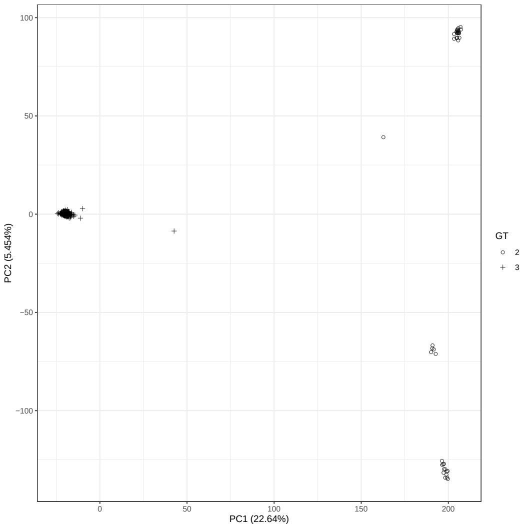


(b) PCA virus data for European samples

Figure S6 Principal component analyses for the human and virus data. a) PCA for infected and uninfected human data of PC1 and PC2. We show in blue 451 individuals of European Ancestry, in orange 74 individuals of Asian ancestry, in light pink 6 individuals of African ancestry from Ansari et al. (2017). From the 1,000 genomes project we add in brown 99 Utah residents with Northern and Western European ancestry (CEU), in black 99 Finnish individuals in Finland (FIN), in green 91 British individuals in England and Scotland (GBR), in pink 107 Spanish individuals (IBS) and in lightgreen 107 Italian individuals (TSI - 107). Proportion of variances for PC1 and PC2 are shown in parentheses. b) PCA for the virus sampled from the 451 European individuals (EUAnsari) from Ansari et al. (2017). Circles refer to HCV genotype 2 samples, while crosses indicate HCV genotype 3 samples.

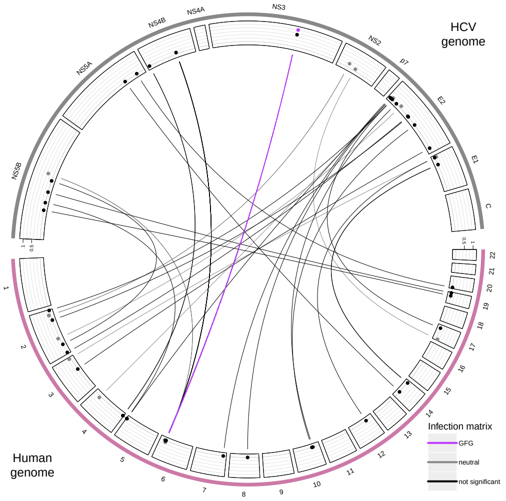


(a) Circos plot for top 200 CSA candidates

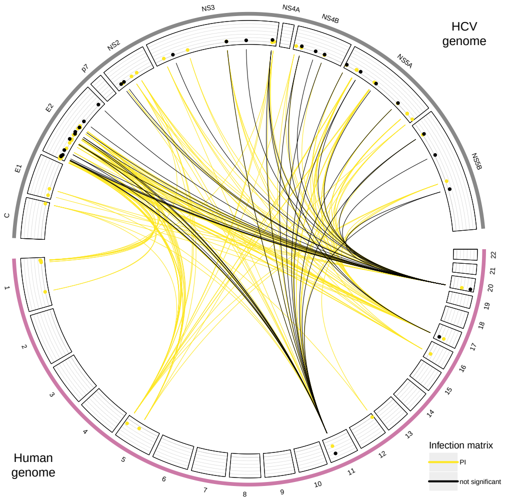


(b) Circos plot for top 200 HP candidates

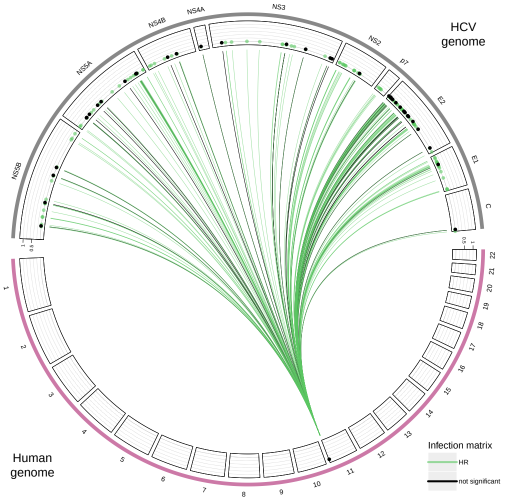


(c) Circos plot for top 200 HS candidates

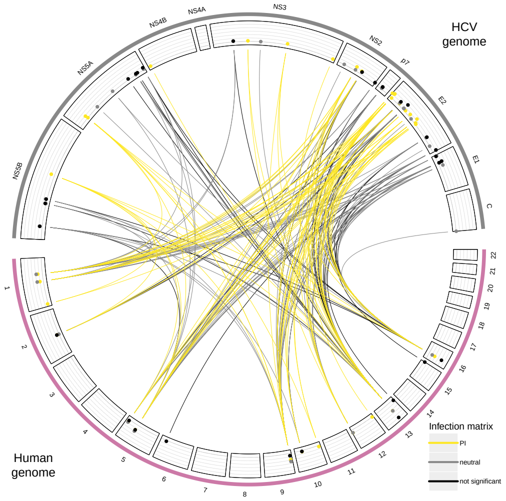


(d) Circos plot for top 200 PI candidates

Figure S7 The genome-to-genome (a.k.a. Circos) plots exhibit the outcomes of the ABC-model choice for the top 200 associations for each index: (a) CSA, (b) HP, (c) HS, and (d) PI. Each plot includes the 200 top associations and their respective infection matrix as inferred by abc model choice. Infection matrices are visible when a single best model was identified (highest overall Bayes factor, Bayes factor compared to all models ¿ 2). All models that had at least one competing model are presented as black lines and are considered not relevant. The inferred matrices are color-coded as follows: purple for $\mathcal{A}_{\mathcal{GFG}}$, green for $\mathcal{A}_{\mathcal{H}}$, yellow for $\mathcal{A}_{\mathcal{P}}$, grey for $\mathcal{A}_{\mathcal{N}}$, and black for not relevant.

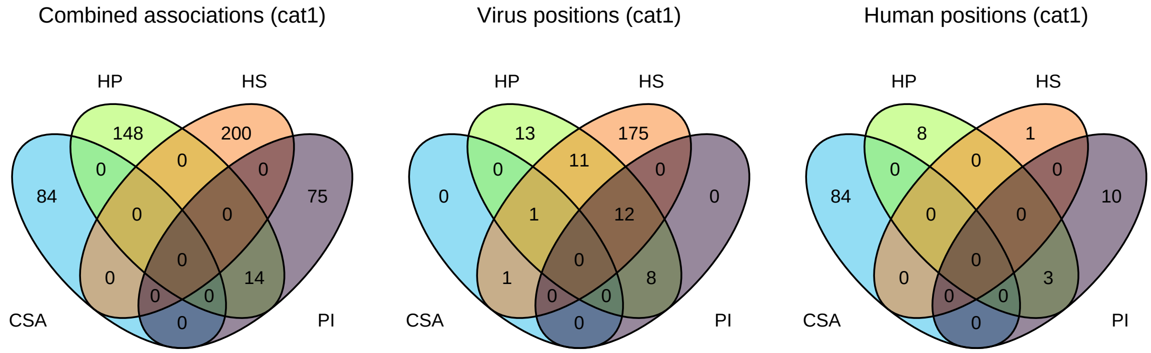


Figure S8 Venn diagram of the overlap of the inferred relevant 535 SNPs-SAAPs associations between indices (CSA, HP, HS, PI). In a) all associations (combination of human and virus position) are shown. All relevant virus sites involved in these associations are shown in b) and all involved human sites are shown in c).

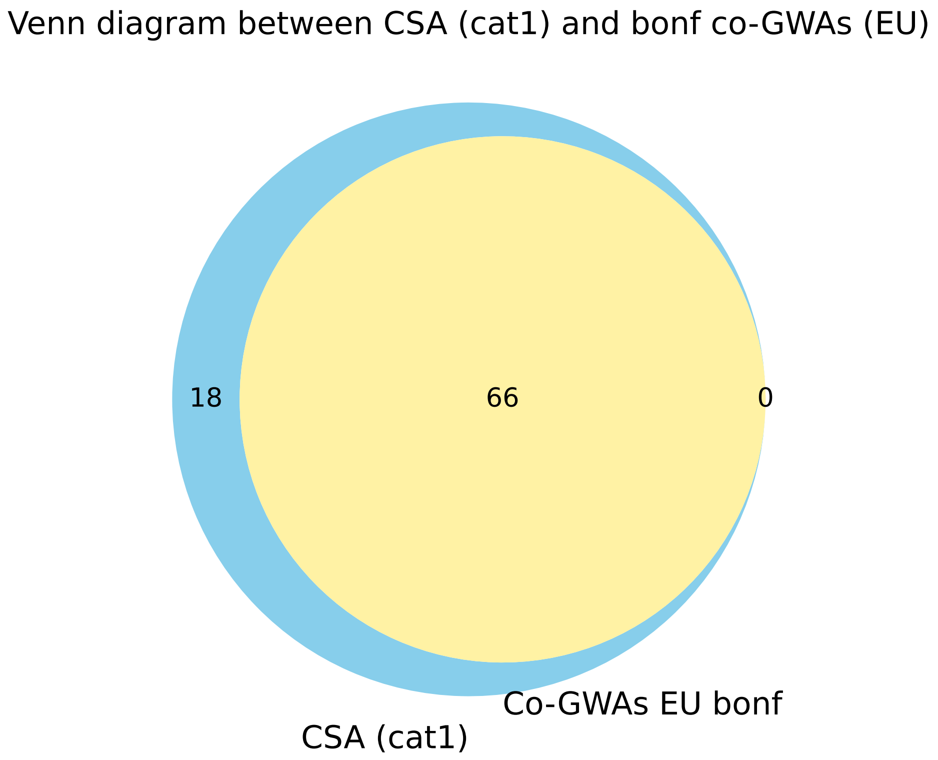


Figure S9  Venn diagram showing the overlap between the relevant 84 candidate associations which have been obtained from the 200 associations with the highest CSA value and the top candidates of our Bonferroni corrected (Bonf p-value=0.05) co-GWAs using the 451 genomes of infected European individuals (and the corresponding HCV strains).

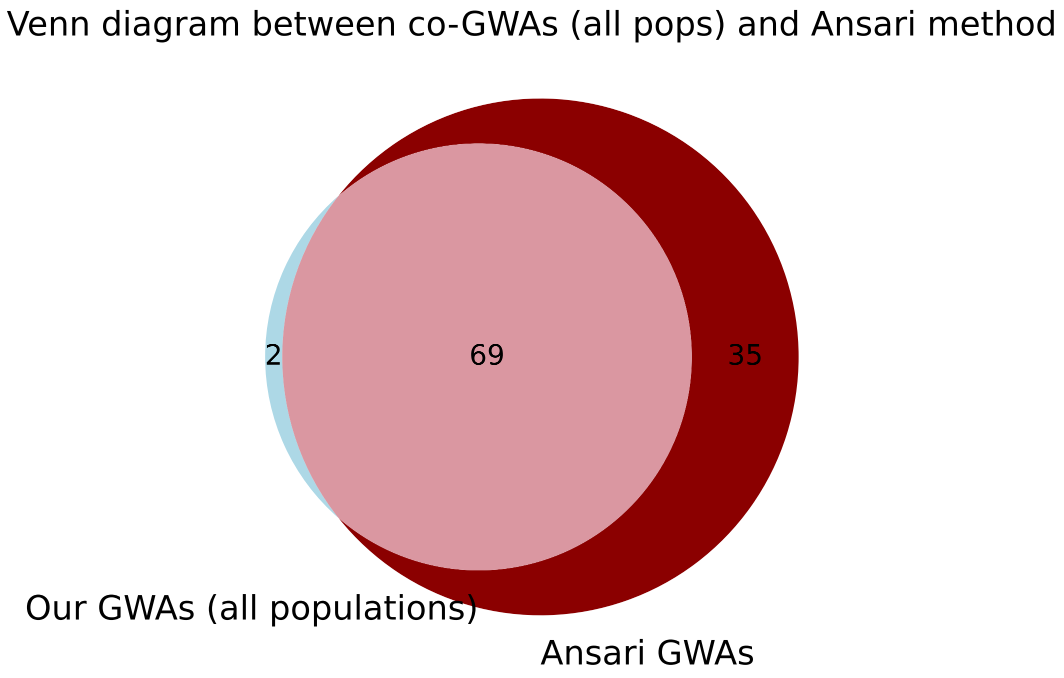


Figure S10 Venn diagram showing the overlap between the top candidates from two co-GWAs containing all 541 individuals from different human populations: the Bonferroni corrected from Ansari et al. (2017) and our Bonferroni corrected one (both with Bonf p-value=0.05).

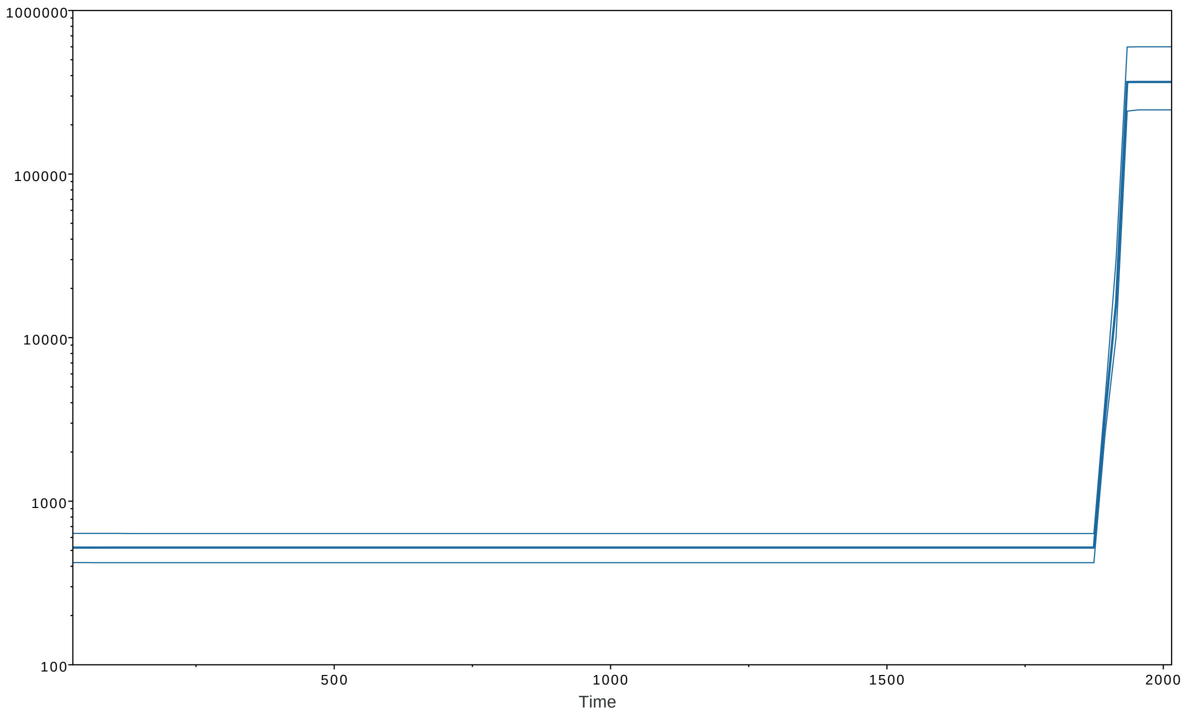


Figure S11 Coalescent Bayesian Skyline plot for the 451 European HCV strains. The darkblue line is the median estimate of the estimated effective population size. The two lightblue lines are the upper and lower bounds of the 95% highest posterior density (HPD) interval. The x-axis is the forward time in years and the y-axis represents the population size on a log-scale.

### Supplementary Tables

Table S1 Expressions for host frequencies in the different infected/uninfected classes.

| Quantity | Description |
|---|---|
| $h_i$ | Frequency of host genotype $i$ in the entire population at the beginning of the generation |
| $p_j$ | Frequency of pathogen genotype $j$ at the beginning of the generation |
| $\alpha_{ij}$ | Probability that pathogen genotype $j$ can infect host genotype $i$ |
| $\phi$ | Proportion of hosts exposed to the pathogen |
| $f_{10} = (1-\phi)h_1$ | frequency of type 1 hosts without pathogen encounter |
| $f_{11} = \phi\alpha_{11}h_1p_1$ | frequency of type 1 hosts infected with pathogens of type 1 |
| $f_{12} = \phi\alpha_{12}h_1p_2$ | frequency of type 1 hosts infected with pathogens of type 2 |
| $f_{13} = \phi\Big[(1-\alpha_{12})\,h_1p_2 + (1-\alpha_{11})\,h_1p_1\Big]$ | frequency of type 1 hosts which had a pathogen encounter but resisted infection |
| $f_{1z} = f_{10} + f_{13}$ | frequency of uninfected genotype 2 hosts in the entire population |
| $f_{20} = (1-\phi)\,h_2$ | frequency of type 2 hosts without pathogen encounter |
| $f_{21} = \phi\alpha_{21}h_2p_1$ | frequency of type 2 hosts infected with pathogens of type 1 |
| $f_{22} = \phi\alpha_{22}h_2p_2$ | frequency of type 2 hosts infected with pathogens of type 2 |
| $f_{23} = \phi\Big[(1-\alpha_{21})\,h_2p_1 + (1-\alpha_{22})\,h_2p_2\Big]$ | frequency of type 2 hosts which had a pathogen encounter but resisted infection |
| $f_{2z} = f_{20} + f_{23}$ | frequency of uninfected genotype 2 hosts in the entire population |
| $\tilde{f} = \sum_{i=1}^{2}\sum_{j=1}^{2} f_{ij}$ | Frequency of **infected** hosts in the population |
| $\tilde{h}_i = \frac{f_{i1} + f_{i2}}{\tilde{f}}$ | Frequency of host genotype $i$ among all **infected** individuals in the population |
| $\tilde{p}_j = \frac{f_{1j} + f_{2j}}{\tilde{f}}$ | Frequency of pathogen genotype $j$ after infections have taken place. |
| $\tilde{f}_{iz} = \frac{f_{iz}}{\sum_{k=1}^{2} f_{kz}}$ | Frequency of hosts of genotype $i$ among all **non-infected** individuals. |
| $\tilde{f}_{ij} = \frac{f_{ij}}{\tilde{f}}$ | Frequency of hosts of genotype $i$ infected by pathogens of genotype $j$ among all **infected** individuals in the population. |

Table S2 Expressions for the various infection/interaction matrices with fully explicit coefficients. Colors indicate if the behaviour of the index with changing pathogen frequency $p_1$. Color coding is as follows: blue = the index is zero, irrespective of the matrix parametrization, green= the index is constant with respect to pathogen frequency, and the constant depends on the parametrization of the matrix, pink= the index changes linearly with pathogen frequency, orange = the relationship between the index and the pathogen frequency is non-linear.

| | HS | PI | CSA[2] | HP |
|---|---|---|---|---|
| $\mathcal{A}_{\mathcal{N}} = \begin{pmatrix} \alpha_{\mathcal{N}} & \alpha_{\mathcal{N}} \\ \alpha_{\mathcal{N}} & \alpha_{\mathcal{N}} \end{pmatrix}$ | 0 | $\lvert\phi^2 \alpha_{\mathcal{N}}^2 (p_2 - p_1)\rvert$ | | $\lvert\phi\alpha_{\mathcal{N}} (1 - \phi\alpha_{\mathcal{N}}) (p_2 - p_1)\rvert$ |
| $\mathcal{A}_{\mathcal{H}} = \begin{pmatrix} \alpha_{\mathcal{H}1} & \alpha_{\mathcal{H}1} \\ \alpha_{\mathcal{H}2} & \alpha_{\mathcal{H}2} \end{pmatrix}$ | $\lvert\phi (\alpha_{\mathcal{H}1} - \alpha_{\mathcal{H}2})\rvert$ | $\lvert\phi^2 \alpha_{\mathcal{H}1}\alpha_{\mathcal{H}2} (p_2 - p_1)\rvert$ | 0 | $= \phi\alpha_{\mathcal{H}1} (1 - \phi\alpha_{\mathcal{H}2})$ $-\phi (\alpha_{\mathcal{H}1} - 2\phi\alpha_{\mathcal{H}1}\alpha_{\mathcal{H}2} + \alpha_{\mathcal{H}2}) p_1$ |
| $\mathcal{A}_{\mathcal{P}} = \begin{pmatrix} \alpha_{\mathcal{P}1} & \alpha_{\mathcal{P}2} \\ \alpha_{\mathcal{P}1} & \alpha_{\mathcal{P}2} \end{pmatrix}$ | 0 | $\lvert\phi^2 (\alpha_{\mathcal{P}2}^2 p_2^2 - \alpha_{\mathcal{P}1}^2 p_1^2)\rvert$ | 0 | $= \phi\alpha_{\mathcal{P}2}(1 - \phi\alpha_{\mathcal{P}2})$ $+\phi\Big[(2\phi\alpha_{\mathcal{P}2}^2 - \alpha_{\mathcal{P}2} - \alpha_{\mathcal{P}1})p_1$ $+ \phi(\alpha_{\mathcal{P}1}^2 - \alpha_{\mathcal{P}2}^2)p_1^2\Big]$ |
| $\mathcal{A}_{\mathcal{MA}} = \begin{pmatrix} \alpha_{\mathcal{M}1} & \alpha_{\mathcal{M}2} \\ \alpha_{\mathcal{M}2} & \alpha_{\mathcal{M}1} \end{pmatrix}$ | $\lvert\phi (\alpha_{\mathcal{M}1} - \alpha_{\mathcal{M}2}) (p_1 - p_2)\rvert$ | $\lvert\phi^2 \alpha_{\mathcal{M}1}\alpha_{\mathcal{M}2}(p_2 - p_1)\rvert$ | $\left\lvert\frac{h_1 h_2 p_1 p 2(\alpha_{\mathcal{M}1}^2 - \alpha_{\mathcal{M}2}^2)^2}{(\alpha_{\mathcal{M}1}p_1 + \alpha_{\mathcal{M}2}p_2)\cdots}\right\rvert$ | $= \phi\alpha_{\mathcal{M}2} (1 - \phi\alpha_{\mathcal{M}1})$ $(p_2 - p_1)$ |
| $\mathcal{A}_{\mathcal{GFG}} = \begin{pmatrix} \alpha_{\mathcal{G}1} & \alpha_{\mathcal{G}1} \\ \alpha_{\mathcal{G}2} & \alpha_{\mathcal{G}1} \end{pmatrix}$ | $\lvert\phi (\alpha_{\mathcal{G}1} - \alpha_{\mathcal{G}2}) p_1\rvert$ | $\lvert\phi^2 \alpha_{\mathcal{G}1} (\alpha_{\mathcal{G}1} p_2^2 - \alpha_{\mathcal{G}2} p_1^2)\rvert$ | $\left\lvert\frac{h_1 h_2 p_1 p_2}{\left(\frac{\alpha_{\mathcal{G}_2}}{(\alpha_{\mathcal{G}_1} - \alpha_{\mathcal{G}_2})} + p_2\right)\left(h_1 + \frac{\alpha_{\mathcal{G}_2}}{(\alpha_{\mathcal{G}_1} - \alpha_{\mathcal{G}_2})}\right)}\right\rvert$ | $= \phi\Big[\alpha_{\mathcal{G}1}p_2 - \alpha_{\mathcal{G}2}p_1$ $-\phi\alpha_{\mathcal{G}1}(\alpha_{\mathcal{G}1}p_2^2 - \alpha_{\mathcal{G}2}p_1^2)\Big]$ |

Table S3  Results of the leave-one-out ABC (rejection) cross-validation for 500 random chosen simulations per infection matrix under low disease encounter rate. For each model 50,000 simulations are produced for $h_1 \sim \mathcal{U}(0.05, 0.5)$, $p_1 \sim \mathcal{U}(0.05, 0.5)$, $\delta = 0.2$, $\phi = 0.05$, $N = 100,000$, $n_{\mathrm{I}} = 902$ haploid and $n_H = 1006$ haploid.

| | Inferred model | | | | | |
|---|---|---|---|---|---|---|
| True model | neutral | GFG | MA | iGFG | HR | PI |
| neutral | 448 | 0 | 0 | 0 | 0 | 52 |
| GFG | 6 | 341 | 33 | 49 | 3 | 68 |
| MA | 0 | 34 | 442 | 24 | 0 | 0 |
| iGFG | 2 | 80 | 112 | 148 | 156 | 2 |
| HR | 0 | 0 | 0 | 8 | 492 | 0 |
| PI | 155 | 64 | 0 | 0 | 0 | 281 |

Table S4  Results of the leave-one-out ABC (rejection) cross-validation for 500 random chosen simulations per infection matrix under low disease encounter rate. For each model 50,000 simulations are produced for $h_1 \sim \mathcal{U}(0.05, 0.5)$, $p_1 \sim \mathcal{U}(0.05, 0.5)$, $\delta = 0.3$, $\phi = 0.05$, $N = 100,000$, $n_{\mathrm{I}} = 902$ haploid and $n_H = 1006$ haploid.

| | Inferred model | | | | | |
|---|---|---|---|---|---|---|
| True model | neutral | GFG | MA | iGFG | HR | PI |
| neutral | 451 | 0 | 0 | 0 | 0 | 49 |
| GFG | 21 | 294 | 58 | 25 | 23 | 79 |
| MA | 1 | 49 | 423 | 25 | 1 | 1 |
| iGFG | 7 | 105 | 129 | 75 | 179 | 5 |
| HR | 0 | 3 | 2 | 15 | 480 | 0 |
| PI | 174 | 91 | 11 | 0 | 0 | 224 |

Table S5   Model choice results for the top-200 candidates for the CSA-index when allowing for different deviations $\delta$ from the matrix in the simulated data. Model choice was run for each candidate association using all simulations for the corresponding $\delta$ value (rejection algorithm, tolerance 0.05). For each association the model with the highest overall Bayes factor was chosen as the best model. The corresponding model was considered the single best model when the Bayes factor compared to any other model was $\geq 2$. All models for which the Bayes factor was $< 2$ are listed as competing models.

| Best model | competing models | $\delta$=0.1 | $\delta$=0.2 | $\delta$=0.3 |
|---|---|---|---|---|
| GFG | neutral,PI | 22 | 6 | 0 |
| neutral | GFG,PI | 4 | 0 | 0 |
| GFG | | 73 | 120 | 113 |
| neutral | GFG | 10 | 0 | 0 |
| GFG | iGFG | 1 | 21 | 70 |
| GFG | neutral | 71 | 2 | 0 |
| neutral | | 9 | 0 | 0 |
| GFG | PI | 10 | 50 | 5 |
| iGFG | GFG | 0 | 1 | 10 |
| GFG | iGFG,PI | 0 | 0 | 1 |
| iGFG | GFG,HR | 0 | 0 | 1 |

Table S6   Model choice results for the top-200 candidates for the HS-index when allowing for different deviations $\delta$ from the matrix in the simulated data. Model choice was run for each candidate association using all simulations for the corresponding $\delta$ value (rejection algorithm, tolerance 0.05). For each association the model with the highest overall Bayes factor was chosen as the best model. The corresponding model was considered the single best model when the Bayes factor compared to any other model was $\geq 2$. All models for which the Bayes factor was $< 2$ are listed as competing models.

| Best model | competing models | $\delta$=0.1 | $\delta$=0.2 | $\delta$=0.3 |
|---|---|---|---|---|
| HR | | 158 | 169 | 200 |
| HR | iGFG | 42 | 31 | 0 |

Table S7 Model choice results for the top-200 candidates for the HP-index when allowing for different deviations $\delta$ from the matrix in the simulated data. Model choice was run for each candidate association using all simulations for the corresponding $\delta$ value (rejection algorithm, tolerance 0.05). For each association the model with the highest overall Bayes factor was chosen as the best model. The corresponding model was considered the single best model when the Bayes factor compared to any other model was $\geq 2$. All models for which the Bayes factor was $< 2$ are listed as competing models.

| Best model | competing models | $\delta$=0.1 | $\delta$=0.2 | $\delta$=0.3 |
|---|---|---|---|---|
| PI | neutral | 34 | 12 | 2 |
| neutral | PI | 4 | 0 | 0 |
| PI | | 162 | 183 | 121 |
| PI | HR, iGFG | 0 | 1 | 52 |
| PI | HR, neutral | 0 | 1 | 0 |
| PI | HR, iGFG, neutral | 0 | 2 | 13 |
| PI | iGFG | 0 | 1 | 6 |
| iGFG | HR, PI | 0 | 0 | 3 |
| HR | iGFG, neutral, PI | 0 | 0 | 2 |
| PI | GFG, HR, iGFG | 0 | 0 | 1 |

Table S8 Model choice results for the top-200 candidates for the PI-index when allowing for different deviations $\delta$ from the matrix in the simulated data. Model choice was run for each candidate association using all simulations for the corresponding $\delta$ value (rejection algorithm, tolerance 0.05). For each association the model with the highest overall Bayes factor was chosen as the best model. The corresponding model was considered the single best model when the Bayes factor compared to any other model was $\geq 2$. All models for which the Bayes factor was $< 2$ are listed as competing models.

| Best model | competing models | $\delta$=0.1 | $\delta$=0.2 | $\delta$=0.3 |
|---|---|---|---|---|
| neutral | | 76 | 38 | 33 |
| neutral | PI | 35 | 72 | 78 |
| PI | | 89 | 89 | 89 |
| PI | neutral | 0 | 1 | 0 |
