## Supplementary material for "Inference of host-pathogen interaction matrices from genome-wide polymorphism data": SItext-methods

### Supplementary Text S1: extended methods

Hanna Märkle[a,b,1], Sona John[a,1], Lukas Metzger[a,1], STOP-HCV Consortium[c], M Azim Ansari[c], Vincent Pedergnana[d], Aurélien Tellier[a,1]

[a]*Population Genetics, Department of Life Science Systems, School of Life Sciences, Technical University of Munich, 85354 Freising, Germany*
[b]*Center for Genomics & Systems Biology, New York University, New York, NY 10003, USA*
[c]*Nuffield Department of Medicine, Peter Medawar Building for Pathogen Research, University of Oxford, Oxford, UK*
[d]*Laboratoire MIVEGEC (UMR CNRS 5290, UR IRD 224, UM), Montpellier, France*

### Theoretical model and analytical derivations

#### Infection matrices of interest and indices values

We derive expressions of our four indices for the five infection matrices in Table S1 to gain an understanding of the behaviour of the proposed indices.

Table S1 Infection matrices of interest. The infection matrix $\mathcal{A}$ determines the outcome of the interaction between host genotypes (rows) and pathogen genotypes (columns). Each $\alpha_{ij}$ can be interpreted either as the probability for a given host to be infected or as the degree of infection (disease severity or partial resistance).

| **neutral infection matrix** $\mathcal{A}_N$ | **differential host resistance** $\mathcal{A}_H$ | **differential pathogen infectivity** $\mathcal{A}_P$ | **simplified GxG interaction** $\mathcal{A}_{GxG}$ | **matching alleles (MA)** $\mathcal{A}_{\mathcal{MA}}$ | **gene-for-gene (GFG and iGFG)** $\mathcal{A}_{\mathcal{GFG}}$ |
|---|---|---|---|---|---|
| $\begin{pmatrix} \alpha_n & \alpha_n \\ \alpha_n & \alpha_n \end{pmatrix}$ | $\begin{pmatrix} \alpha_{\mathcal{H}1} & \alpha_{\mathcal{H}1} \\ \alpha_{\mathcal{H}2} & \alpha_{\mathcal{H}2} \end{pmatrix}$ | $\begin{pmatrix} \alpha_{\mathcal{P}1} & \alpha_{\mathcal{P}2} \\ \alpha_{\mathcal{P}1} & \alpha_{\mathcal{P}2} \end{pmatrix}$ | $\begin{pmatrix} 1 & \alpha_{12} \\ \alpha_{21} & 1 \end{pmatrix}$ | $\begin{pmatrix} \alpha_{\mathcal{M}1} & \alpha_{\mathcal{M}2} \\ \alpha_{\mathcal{M}2} & \alpha_{\mathcal{M}1} \end{pmatrix}$ | $\begin{pmatrix} \alpha_{\mathcal{G}1} & \alpha_{\mathcal{G}1} \\ \alpha_{\mathcal{G}2} & \alpha_{\mathcal{G}1} \end{pmatrix}$ |

*The neutral infection matrix ($\mathcal{A}_N$)* characterizes associations between host and pathogen loci not involved into the interaction and is defined by $\alpha_{ij} = \alpha_n, \forall i, j$ and $0 \leq \alpha_n \leq 1$.
*The differential host resistance matrix ($\mathcal{A}_H$)* is defined by two host genotypes having different levels of resistance against any pathogen genotype ($\alpha_{1j} = \alpha_{\mathcal{H}1}$ and $\alpha_{2j} = \alpha_{\mathcal{H}2} \quad, \forall j$). In the extreme case there is a universally resistant host genotype $k$ ($\alpha_{\mathcal{H}k} = 0$) and a universally susceptible host genotype $l$ ($\alpha_{\mathcal{H}l} = 1$).
Similarly, *the differential pathogen infectivity matrix* ($\mathcal{A}_P$) is characterized by two pathogen genotypes with overall differences in infection capability on either host genotype ($\alpha_{i1} = \alpha_{\mathcal{P}1}$ and $\alpha_{i2} = \alpha_{\mathcal{P}2} \quad, \forall i$ ). In the most extreme case there is a universally infective pathogen genotype $x$ ($\alpha_{\mathcal{P}x} = 1$) and a universally non-infective pathogen genotype $y$ ($\alpha_{\mathcal{P}y} = 0$).
In a *matching-alleles (inverse matching-alleles interaction) ($\mathcal{A}_{\mathcal{MA}}$)* pathogen genotypes are better (worse) in infecting matching (non-matching) host genotypes than in infecting non-matching (matching) host genotypes ($\alpha_{ii} = \alpha_{\mathcal{M}1}$ and $\alpha_{ij} = \alpha_{\mathcal{M}2}$ for $i \neq j$). For a matching-alleles matrix $\alpha_{\mathcal{M}1} > \alpha_{\mathcal{M}2}$ and for an inverse matching alleles matrix

$\alpha_{\mathcal{M}1} < \alpha_{\mathcal{M}2}$.

*Gene-for-gene/inverse gene-for-gene interactions* are characterized by differences in susceptibility of the host genotypes and differences in infectivity of the two pathogen genotypes. In a perfect gene-for-gene matrix there is universally susceptible host genotype $l$ ($\alpha_{lj} = 1 \quad, \forall j$) and universally infective pathogen genotype $x$ ($\alpha_{ix} = 1 \quad, \forall i$). In a perfect inverse gene-for-gene interaction there is universally resistant host genotype $l$ ($\alpha_{lj} = 0 \quad, \forall j$) and a universally non-infective pathogen genotype $x$ ($\alpha_{ix} = 0 \quad, \forall i$).

Here, we derive the values of the indices for various infection matrices presented in Table S1 and discuss the general behaviour of each index.

*CSA index*

$$
\begin{aligned}
\mathrm{CSA}^2_{\mathcal{A}_\mathcal{N}} &= \left| \frac{h_1 h_2 p_1 p_2 \left(\alpha_{11}\alpha_{22} - \alpha_{12}\alpha_{21}\right)^2}{\left(\alpha_{11}p_1 + \alpha_{12}p_2\right)\left(\alpha_{21}p_1 + \alpha_{22}p_2\right)\left(\alpha_{11}h_1 + \alpha_{21}h_2\right)\left(\alpha_{12}h_1 + \alpha_{22}h_2\right)} \right|, \\
&= \left| \frac{h_1 h_2 p_1 p_2 \left(\alpha_\mathcal{N}^2 - \alpha_\mathcal{N}^2\right)^2}{\alpha_\mathcal{N}\alpha_\mathcal{N}\alpha_\mathcal{N}\alpha_\mathcal{N}} \right|, \\
&= 0
\end{aligned} \tag{1}
$$

$$
\begin{aligned}
\mathrm{CSA}^2_{\mathcal{A}_\mathcal{H}} &= \left| \frac{h_1 h_2 p_1 p_2 \left(\alpha_{11}\alpha_{22} - \alpha_{12}\alpha_{21}\right)^2}{\left(\alpha_{11}p_1 + \alpha_{12}p_2\right)\left(\alpha_{21}p_1 + \alpha_{22}p_2\right)\left(\alpha_{11}h_1 + \alpha_{21}h_2\right)\left(\alpha_{12}h_1 + \alpha_{22}h_2\right)} \right|, \\
&= \left| \frac{h_1 h_2 p_1 p_2 \left(\alpha_{\mathcal{H}_1}\alpha_{\mathcal{H}_2} - \alpha_{\mathcal{H}_1}\alpha_{\mathcal{H}_2}\right)^2}{\alpha_{\mathcal{H}_1}\alpha_{\mathcal{H}_2}\left(\alpha_{\mathcal{H}_1}h_1 + \alpha_{\mathcal{H}_2}h_2\right)^2} \right| \\
&= 0
\end{aligned} \tag{2}
$$

$$
\begin{aligned}
\mathrm{CSA}^2_{\mathcal{A}_\mathcal{P}} &= \left| \frac{h_1 h_2 p_1 p_2 \left(\alpha_{11}\alpha_{22} - \alpha_{12}\alpha_{21}\right)^2}{\left(\alpha_{11}p_1 + \alpha_{12}p_2\right)\left(\alpha_{21}p_1 + \alpha_{22}p_2\right)\left(\alpha_{11}h_1 + \alpha_{21}h_2\right)\left(\alpha_{12}h_1 + \alpha_{22}h_2\right)} \right|, \\
&= \left| \frac{h_1 h_2 p_1 p_2 \left(\alpha_{\mathcal{P}_1}\alpha_{\mathcal{P}_2} - \alpha_{\mathcal{P}_2}\alpha_{\mathcal{P}_1}\right)^2}{\left(\alpha_{\mathcal{P}_1}p_1 + \alpha_{\mathcal{P}_2}p_2\right)^2\alpha_{\mathcal{P}_1}\alpha_{\mathcal{P}_2}} \right| \\
&= 0
\end{aligned} \tag{3}
$$

$$\mathrm{CSA}^2_{\mathcal{AMA}} = \left|\frac{h_1h_2p_1p_2\left(\alpha_{11}\alpha_{22}-\alpha_{12}\alpha_{21}\right)^2}{\left(\alpha_{11}p_1+\alpha_{12}p_2\right)\left(\alpha_{21}p_1+\alpha_{22}p_2\right)\left(\alpha_{11}h_1+\alpha_{21}h_2\right)\left(\alpha_{12}h_1+\alpha_{22}h_2\right)}\right|, \quad (4)$$

$$= \left|\frac{h_1h_2p_1p_2\left(\alpha^2_{\mathcal{M}_1}-\alpha^2_{\mathcal{M}_2}\right)^2}{\left(\alpha_{\mathcal{M}_1}p_1+\alpha_{\mathcal{M}_2}p_2\right)\left(\alpha_{\mathcal{M}_2}p_1+\alpha_{\mathcal{M}_1}p_2\right)\left(\alpha_{\mathcal{M}_1}h_1+\alpha_{\mathcal{M}_2}h_2\right)\left(\alpha_{\mathcal{M}_2}h_1+\alpha_{\mathcal{M}_1}h_2\right)}\right|$$

Note the following equations require $\alpha_{\mathcal{G}1} > 0$. CSA is not defined if $\alpha_{\mathcal{G}1} = 0$ as, in this case, one pathogen genotype cannot infect.

$$\mathrm{CSA}^2_{\mathcal{GFG}} = \left|\frac{h_1h_2p_1p_2\left(\alpha_{11}\alpha_{22}-\alpha_{12}\alpha_{21}\right)^2}{\left(\alpha_{11}p_1+\alpha_{12}p_2\right)\left(\alpha_{21}p_1+\alpha_{22}p_2\right)\left(\alpha_{11}h_1+\alpha_{21}h_2\right)\left(\alpha_{12}h_1+\alpha_{22}h_2\right)}\right|, \quad (5)$$

$$= \left|\frac{h_1h_2p_1p_2\left(\alpha_{\mathcal{G}_1}\alpha_{\mathcal{G}_1}-\alpha_{\mathcal{G}_1}\alpha_{\mathcal{G}_2}\right)^2}{\left(\alpha_{\mathcal{G}_1}p_1+\alpha_{\mathcal{G}_1}p_2\right)\left(\alpha_{\mathcal{G}_2}p_1+\alpha_{\mathcal{G}_1}p_2\right)\left(\alpha_{\mathcal{G}_1}h_1+\alpha_{\mathcal{G}_2}h_2\right)\left(\alpha_{\mathcal{G}_1}h_1+\alpha_{\mathcal{G}_1}h_2\right)}\right|$$

$$= \left|\frac{h_1h_2p_1p_2\left(\alpha_{\mathcal{G}_1}\left(\alpha_{\mathcal{G}_1}-\alpha_{\mathcal{G}_2}\right)\right)^2}{\alpha_{\mathcal{G}_1}\left(\alpha_{\mathcal{G}_2}p_1+\alpha_{\mathcal{G}_1}p_2\right)\left(\alpha_{\mathcal{G}_1}h_1+\alpha_{\mathcal{G}_2}h_2\right)\alpha_{\mathcal{G}_1}}\right|$$

$$= \left|\frac{h_1h_2p_1p_2\alpha^2_{\mathcal{G}_1}\left(\alpha_{\mathcal{G}_1}-\alpha_{\mathcal{G}_2}\right)^2}{\alpha_{\mathcal{G}_1}\left(\alpha_{\mathcal{G}_2}p_1+\alpha_{\mathcal{G}_1}p_2\right)\left(\alpha_{\mathcal{G}_1}h_1+\alpha_{\mathcal{G}_2}h_2\right)\alpha_{\mathcal{G}_1}}\right|$$

$$= \left|\frac{h_1h_2p_1p_2\left(\alpha_{\mathcal{G}_1}-\alpha_{\mathcal{G}_2}\right)^2}{\left(\alpha_{\mathcal{G}_2}p_1+\alpha_{\mathcal{G}_1}p_2\right)\left(\alpha_{\mathcal{G}_1}h_1+\alpha_{\mathcal{G}_2}h_2\right)}\right|$$

Note the following simplifications are only valid, if and only if, $\alpha_{\mathcal{G}1} \neq \alpha_{\mathcal{G}2}$ and $\alpha_{\mathcal{G}1} > 0$.

$$\begin{aligned}
\mathrm{CSA}^2_{\mathcal{GFG}} &= \left| \frac{h_1 h_2 p_1 p_2 \left(\alpha_{\mathcal{G}_1} - \alpha_{\mathcal{G}_2}\right)^2}{\left(\alpha_{\mathcal{G}_2} p_1 + \alpha_{\mathcal{G}_1} p_2\right)\left(\alpha_{\mathcal{G}_1} h_1 + \alpha_{\mathcal{G}_2} h_2\right)} \right| \\
&= \left| \frac{h_1 h_2 p_1 p_2 \left(\alpha_{\mathcal{G}_1} - \alpha_{\mathcal{G}_2}\right)^2}{\left(\alpha_{\mathcal{G}_2}\left(1 - p_2\right) + \alpha_{\mathcal{G}_1} p_2\right)\left(\alpha_{\mathcal{G}_1} h_1 + \alpha_{\mathcal{G}_2}\left(1 - h_1\right)\right)} \right| \\
&= \left| \frac{h_1 h_2 p_1 p_2 \left(\alpha_{\mathcal{G}_1} - \alpha_{\mathcal{G}_2}\right)^2}{\left(\alpha_{\mathcal{G}_2} - \alpha_{\mathcal{G}_2} p_2 + \alpha_{\mathcal{G}_1} p_2\right)\left(\alpha_{\mathcal{G}_1} h_1 + \alpha_{\mathcal{G}_2} - \alpha_{\mathcal{G}_2} h_1\right)} \right| \\
&= \left| \frac{h_1 h_2 p_1 p_2 \left(\alpha_{\mathcal{G}_1} - \alpha_{\mathcal{G}_2}\right)^2}{\left(\left(\alpha_{\mathcal{G}_1} - \alpha_{\mathcal{G}_2}\right)\frac{\alpha_{\mathcal{G}_2}}{\left(\alpha_{\mathcal{G}_1} - \alpha_{\mathcal{G}_2}\right)} + \left(\alpha_{\mathcal{G}_1} - \alpha_{\mathcal{G}_2}\right) p_2\right)\left(\left(\alpha_{\mathcal{G}_1} - \alpha_{\mathcal{G}_2}\right) h_1 + \left(\alpha_{\mathcal{G}_1} - \alpha_{\mathcal{G}_2}\right)\frac{\alpha_{\mathcal{G}_2}}{\left(\alpha_{\mathcal{G}_1} - \alpha_{\mathcal{G}_2}\right)}\right)} \right| \\
&= \left| \frac{h_1 h_2 p_1 p_2 \left(\alpha_{\mathcal{G}_1} - \alpha_{\mathcal{G}_2}\right)^2}{\left(\alpha_{\mathcal{G}_1} - \alpha_{\mathcal{G}_2}\right)\left(\frac{\alpha_{\mathcal{G}_2}}{\left(\alpha_{\mathcal{G}_1} - \alpha_{\mathcal{G}_2}\right)} + p_2\right)\left(\alpha_{\mathcal{G}_1} - \alpha_{\mathcal{G}_2}\right)\left(h_1 + \frac{\alpha_{\mathcal{G}_2}}{\left(\alpha_{\mathcal{G}_1} - \alpha_{\mathcal{G}_2}\right)}\right)} \right| \\
&= \left| \frac{h_1 h_2 p_1 p_2}{\left(\frac{\alpha_{\mathcal{G}_2}}{\left(\alpha_{\mathcal{G}_1} - \alpha_{\mathcal{G}_2}\right)} + p_2\right)\left(h_1 + \frac{\alpha_{\mathcal{G}_2}}{\left(\alpha_{\mathcal{G}_1} - \alpha_{\mathcal{G}_2}\right)}\right)} \right|
\end{aligned} \tag{6}$$

The value of the CSA index is always zero irrespective of pathogen frequencies and $\phi$ for a neutral matrix, any universally resistant host genotype matrix ($\mathcal{A}_H$) and any universally infective pathogen matrix $\mathcal{A}_P$. For arbitrary matching-alleles (MA) matrices and gene-for-gene (GFG, iGFG) matrices the normalized index scales non-linearly with pathogen frequencies. For given pathogen frequencies the index is indistinguishable for a perfect matching alleles matrix ($\alpha_{\mathcal{M}1} = 1$, $\alpha_{\mathcal{M}2} = 0$) and a perfect gene-for-gene matrix ($\alpha_{\mathcal{G}1} = 1$, $\alpha_{\mathcal{G}2} = 0$). For a perfect inverse gene-for-gene matrix ($\alpha_{\mathcal{G}1} = 0$, $\alpha_{\mathcal{G}2} = 1$) the index becomes zero irrespective of pathogen frequencies.

*HS index*

$$\begin{aligned}
\mathrm{HS}_{\mathcal{A}_\mathcal{N}} &= \left|\phi\Big[\left(\alpha_{11} - \alpha_{21}\right) p_1 + \left(\alpha_{12} - \alpha_{22}\right) p_2\Big]\right| \\
&= \left|\phi\Big[\left(\alpha_\mathcal{N} - \alpha_\mathcal{N}\right) p_1 + \left(\alpha_\mathcal{N} - \alpha_\mathcal{N}\right) p_2\Big]\right| \\
&= 0
\end{aligned} \tag{7}$$

$$
\begin{aligned}
\mathrm{HS}_{\mathcal{A}_{\mathcal{H}}} &= \left|\phi\Big[(\alpha_{11}-\alpha_{21})\,p_1+(\alpha_{12}-\alpha_{22})\,p_2\Big]\right| \\
&= |\phi\,(\alpha_{\mathcal{H}1}-\alpha_{\mathcal{H}2})\,(p_1+p_2)| \\
&= |\phi\,(\alpha_{\mathcal{H}1}-\alpha_{\mathcal{H}2})|
\end{aligned} \tag{8}
$$

$$
\begin{aligned}
\mathrm{HS}_{\mathcal{A}_{\mathcal{P}}} &= \left|\phi\Big[(\alpha_{11}-\alpha_{21})\,p_1+(\alpha_{12}-\alpha_{22})\,p_2\Big]\right| \\
&= \left|\phi\Big[(\alpha_{\mathcal{P}1}-\alpha_{\mathcal{P}1})\,p_1+(\alpha_{\mathcal{P}2}-\alpha_{\mathcal{P}2})\,p_2\Big]\right| \\
&= 0
\end{aligned} \tag{9}
$$

$$
\begin{aligned}
\mathrm{HS}_{\mathcal{A}_{\mathcal{MA}}} &= \left|\phi\Big[(\alpha_{11}-\alpha_{21})\,p_1+(\alpha_{12}-\alpha_{22})\,p_2\Big]\right| \\
&= \left|\phi\Big[(\alpha_{\mathcal{M}1}-\alpha_{\mathcal{M}2})\,p_1+(\alpha_{\mathcal{M}2}-\alpha_{\mathcal{M}1})\,p_2\Big]\right| \\
&= \left|\phi\Big[(\alpha_{\mathcal{M}1}-\alpha_{\mathcal{M}2})\,(p_1-p_2)\Big]\right|
\end{aligned} \tag{10}
$$

$$
\begin{aligned}
\mathrm{HS}_{\mathcal{A}_{\mathcal{GFG}}} &= \left|\phi\Big[(\alpha_{11}-\alpha_{21})\,p_1+(\alpha_{12}-\alpha_{22})\,p_2\Big]\right| \\
&= \left|\phi\Big[(\alpha_{\mathcal{G}1}-\alpha_{\mathcal{G}2})\,p_1+(\alpha_{\mathcal{G}1}-\alpha_{\mathcal{G}1})\,p_2\Big]\right| \\
&= |\phi\,(\alpha_{\mathcal{G}1}-\alpha_{\mathcal{G}2})\,p_1|
\end{aligned} \tag{11}
$$

HS is always zero for a neutral infection matrix irrespective of pathogen frequencies and $\alpha_{\mathcal{N}}$. For the universally resistant host matrix $\mathcal{A}_{\mathcal{H}}$ the value of the index is independent of the pathogen frequencies and is equal to $\phi \cdot (\alpha_{\mathcal{H}1} - \alpha_{\mathcal{H}2})$ (see Eq. 8).
For the 'universally infective pathogen' matrix ($\alpha_{\mathcal{N}}$) the index value is 0 irrespective of host frequencies, pathogen frequencies and $\phi$ (see Eq. 9 for the calculation).
For any generalized (but still symmetric matching-alleles matrix including the 'perfect' matching-alleles matrix) $\mathcal{A}_{\mathcal{MA}}$ the index value is proportional to $p_1 - p_2$ and the scaling

factor is $\phi(\alpha_{\mathcal{M}1} - \alpha_{\mathcal{M}2})$ (see Eq. 10 ).

For any gene-for gene matrix $\mathcal{A}_{\mathcal{GFG}}$ the index value is proportional to $p_1$ and the scaling factor is $\phi(\alpha_{\mathcal{G}1} - \alpha_{\mathcal{G}2})$ (see Eq. 11).

*PI index*

$$\begin{aligned} \mathrm{PI}_{\mathcal{A}_{\mathcal{N}}} &= \left|\phi^2\left(p_2^2\alpha_{12}\alpha_{22} - p_1^2\alpha_{11}\alpha_{21}\right)\right| \\ &= \left|\phi^2\left(p_2^2\alpha_{\mathcal{N}}^2 - p_1^2\alpha_{\mathcal{N}}^2\right)\right| \\ &= \left|\phi^2\alpha_{\mathcal{N}}^2\left(p_2^2 - p_1^2\right)\right| \\ &= \left|\phi^2\alpha_{\mathcal{N}}^2\left((1-p_1)^2 - p_1^2\right)\right| \\ &= \left|\phi^2\alpha_{\mathcal{N}}^2\left(1 - 2p_1 + p_1^2 - p_1^2\right)\right| \\ &= \left|\phi^2\alpha_{\mathcal{N}}^2\left(p_1 + p_2 - 2p_1\right)\right| \\ &= \left|\phi^2\alpha_{\mathcal{N}}^2\left(p_2 - p_1\right)\right| \end{aligned} \tag{12}$$

$$\begin{aligned} \mathrm{PI}_{\mathcal{A}_{\mathcal{H}}} &= \left|\phi^2\left(p_2^2\alpha_{12}\alpha_{22} - p_1^2\alpha_{11}\alpha_{21}\right)\right| \\ &= \left|\phi^2\left(p_2^2\alpha_{\mathcal{H}1}\alpha_{\mathcal{H}2} - p_1^2\alpha_{\mathcal{H}1}\alpha_{\mathcal{H}2}\right)\right| \\ &= \left|\phi^2\alpha_{\mathcal{H}1}\alpha_{\mathcal{H}2}\left(p_2 - p_1\right)\right| \end{aligned} \tag{13}$$

$$\begin{aligned} \mathrm{PI}_{\mathcal{A}_{\mathcal{P}}} &= \left|\phi^2\left(p_2^2\alpha_{12}\alpha_{22} - p_1^2\alpha_{11}\alpha_{21}\right)\right| \\ &= \left|\phi^2\left(p_2^2\alpha_{\mathcal{P}2}\alpha_{\mathcal{P}2} - p_1^2\alpha_{\mathcal{P}1}\alpha_{\mathcal{P}1}\right)\right| \\ &= \left|\phi^2\left(p_2\alpha_{\mathcal{P}2} - p_1\alpha_{\mathcal{P}1}\right)\left(p_2\alpha_{\mathcal{P}2} + p_1\alpha_{\mathcal{P}1}\right)\right| \end{aligned} \tag{14}$$

$$\begin{aligned} \mathrm{PI}_{\mathcal{A}_{\mathcal{MA}}} &= \left|\phi^2\left(p_2^2\alpha_{12}\alpha_{22} - p_1^2\alpha_{11}\alpha_{21}\right)\right| \\ &= \left|\phi^2\left(p_2^2\alpha_{\mathcal{M}2}\alpha_{\mathcal{M}1} - p_1^2\alpha_{\mathcal{M}1}\alpha_{\mathcal{M}2}\right)\right| \\ &= \left|\phi^2\alpha_{\mathcal{M}1}\alpha_{\mathcal{M}2}\left(p_2 - p_1\right)\right| \end{aligned} \tag{15}$$

$$\begin{aligned}
\mathrm{PI}_{\mathcal{A}_{\mathcal{GFG}}} &= \left|\phi^2\left(p_2^2\alpha_{12}\alpha_{22} - p_1^2\alpha_{11}\alpha_{21}\right)\right| \\
&= \left|\phi^2\left(p_2^2\alpha_{\mathcal{G}1}\alpha_{\mathcal{G}1} - p_1^2\alpha_{\mathcal{G}1}\alpha_{\mathcal{G}2}\right)\right| \\
&= \left|\phi^2\alpha_{\mathcal{G}1}\left(\alpha_{\mathcal{G}1}p_2^2 - p_1^2\alpha_{\mathcal{G}2}\right)\right|
\end{aligned} \tag{16}$$

For the universally resistant host matrix $\mathcal{A}_H$ the index is proportional to $p_2 - p_1$ and the scaling factor is $\phi^2\alpha_{\mathcal{H}1}\alpha_{\mathcal{H}2}$.

PI exhibits a non-linear relationship with pathogen frequencies for the universally infective pathogen matrix ($\mathcal{A}_P$). If for either pathogen genotype $k$ all entries of the infection matrix are $\alpha_{\mathcal{P}k} \approx 0$ the index is proportional to the square of the frequency of the other pathogen genotype $p_{j\neq k}^2$. For the generalized matching-alleles matrix the index is proportional to $p_2 - p_1$ and scales with $\phi^2\alpha_{\mathcal{M}1}\alpha_{\mathcal{M}2}$ (see Eq. 15). Accordingly, the index will be always zero for a perfect matching-alleles model ($\alpha_{\mathcal{M}1} = 0$, $\alpha_{\mathcal{M}2} = 1$ or $\alpha_{\mathcal{M}1} = 1$, $\alpha_{\mathcal{M}2} = 0$). For an arbitrary GFG-matrix the index scales non-linearly with pathogen frequencies. For a perfect GFG-matrix ($\alpha_{\mathcal{M}1} = 0$, $\alpha_{\mathcal{M}2} = 1$ or $\alpha_{\mathcal{G}1} = 1$, $\alpha_{\mathcal{G}2} = 0$ the index scales linearly with $p_2^2$). For a perfect inverse gene-for-gene model the index becomes 0.

*HP index*

$$\begin{aligned}
\mathrm{HP}_{\mathcal{A}_{\mathcal{N}}} &= \left|\phi\big(\alpha_{12}p_2(1-\phi\alpha_{22}p_2) - \alpha_{21}p_1(1-\phi\alpha_{11}p_1)\big)\right| \\
&= \left|\phi\big(\alpha_{\mathcal{N}}p_2(1-\phi\alpha_{\mathcal{N}}p_2) - \alpha_{\mathcal{N}}p_1(1-\phi\alpha_{\mathcal{N}}p_1)\big)\right| \\
&= |\phi\alpha_{\mathcal{N}}(1-\phi\alpha_{\mathcal{N}})(p_2-p_1)|
\end{aligned} \tag{17}$$

$$\begin{aligned}
\mathrm{HP}_{\mathcal{A}_{\mathcal{H}}} &= \left|\phi\big(\alpha_{12}p_2(1-\phi\alpha_{22}p_2) - \alpha_{21}p_1(1-\phi\alpha_{11}p_1)\big)\right| \\
&= \left|\phi\big(\alpha_{\mathcal{H}1}p_2(1-\phi\alpha_{\mathcal{H}2}p_2) - \alpha_{\mathcal{H}2}p_1(1-\phi\alpha_{\mathcal{H}1}p_1)\big)\right| \\
&= |\phi\alpha_{\mathcal{H}1}(1-\phi\alpha_{\mathcal{H}2}) - \phi(\alpha_{\mathcal{H}1} - 2\phi\alpha_{\mathcal{H}1}\alpha_{\mathcal{H}2} + \alpha_{\mathcal{H}2})p_1|
\end{aligned} \tag{18}$$

$$
\begin{aligned}
\text{HP}_{\mathcal{A}_{\mathcal{P}}} &= \left|\phi\Big[\alpha_{12}p_2(1-\phi\alpha_{22}p_2)-\alpha_{21}p_1(1-\phi\alpha_{11}p_1)\Big]\right| \\
&= \left|\phi\Big[\alpha_{\mathcal{P}2}p_2(1-\phi\alpha_{\mathcal{P}2}p_2)-\alpha_{\mathcal{P}1}p_1(1-\phi\alpha_{\mathcal{P}1}p_1)\Big]\right| \\
&= \left|\phi\alpha_{\mathcal{P}2}(1-\phi\alpha_{\mathcal{P}2})+\phi\Big[(2\phi\alpha_{\mathcal{P}2}^2-\alpha_{\mathcal{P}2}-\alpha_{\mathcal{P}1})p_1+\phi(\alpha_{\mathcal{P}1}^2-\alpha_{\mathcal{P}2}^2)p_1^2\Big]\right|
\end{aligned}
\tag{19}
$$

$$
\begin{aligned}
\text{HP}_{\mathcal{A}_{\mathcal{MA}}} &= \left|\phi\Big[\alpha_{12}p_2(1-\phi\alpha_{22}p_2)-\alpha_{21}p_1(1-\phi\alpha_{11}p_1)\Big]\right| \\
&= \left|\phi\Big[\alpha_{\mathcal{M}2}p_2(1-\phi\alpha_{\mathcal{M}1}p_2)-\alpha_{\mathcal{M}2}p_1(1-\phi\alpha_{\mathcal{M}1}p_1)\Big]\right| \\
&= \left|\phi\Big[\alpha_{\mathcal{M}2}(1-\phi\alpha_{\mathcal{M}1})(p_2-p_1)\Big]\right|
\end{aligned}
\tag{20}
$$

$$
\begin{aligned}
\text{HP}_{\mathcal{A}_{\mathcal{GFG}}} &= \left|\phi\Big[\alpha_{12}p_2(1-\phi\alpha_{22}p_2)-\alpha_{21}p_1(1-\phi\alpha_{11}p_1)\Big]\right| \\
&= \left|\phi\Big[\alpha_{\mathcal{G}1}p_2(1-\phi\alpha_{\mathcal{G}1}p_2)-\alpha_{\mathcal{G}2}p_1(1-\phi\alpha_{\mathcal{G}1}p_1)\Big]\right| \\
&= \left|\phi\Big[\alpha_{\mathcal{G}1}p_2-\alpha_{\mathcal{G}2}p_1-\phi\alpha_{\mathcal{G}1}(\alpha_{\mathcal{G}1}p_2^2-\alpha_{\mathcal{G}2}p_1^2)\Big]\right|
\end{aligned}
\tag{21}
$$

For the neutral matrix the index scales linearly with pathogen frequency. It becomes independent of pathogen frequencies if either $\phi = \alpha_{\mathcal{N}} = 1$ or $\phi = \alpha_{\mathcal{N}} = 0$. For the universally resistant host matrix ($\mathcal{A}_H$) the normalized host partitioning index is proportional to $p_1$. For a perfect universally resistant host interaction matrix ($\alpha_{\mathcal{H}1} = 1$ and $\alpha_{\mathcal{H}2} = 0$ or $\alpha_{\mathcal{H}1} = 0$ and $\alpha_{\mathcal{H}2} = 1$) the index is inversely proportional to $p_1$ with a scaling factor of $\phi$ and the intercept being $0$.

The relationship between the HP-index and pathogen frequencies is non-linear for a generalized universally infective pathogen matrix the HP-index. For a perfect universally infective pathogen matrix with $\alpha_{\mathcal{P}1} = 1$ and $\alpha_{\mathcal{P}2} = 0$) the index simplifies to $-\phi p_1^2(1-\phi p_1)$. Hence, the index monotonically decreases with increasing frequency of pathogen genotype 1 if $\phi \leq 0.5$. For a matching-alleles matrix the index scales linearly with pathogen frequencies.

For a perfect matching-alleles matrix ($\alpha_{\mathcal{M}1} = 1$ and $\alpha_{\mathcal{M}2} = 0$) the index becomes zero. For a perfect inverse-matching alleles matrix the index scales linearly with pathogen frequencies and the scaling factor is $\phi$. For a general gene-for-gene matrix the index scales non-linearly with pathogen frequencies. The index becomes zero for a perfect inverse-gene-for-gene matrix ($\alpha_{\mathcal{G}1} = 0$ and $\alpha_{\mathcal{G}2} = 1$)

### ABC inference method

#### Generating simulations for the ABC model choice

We simulated $m = 50,000$ datasets for each matrix of interest. Simulations were implemented using the following recipe:

1. Draw a random value for $h_1$ from $\mathcal{U}(0.05, 0.5)$
2. Draw a random value for $p_1$ from $\mathcal{U}(0.05, 0.5)$
3. Randomly draw one valid matrix configuration for the matrix of interest (see Table S2 for each matrix).
4. Randomly draw the coefficients of the matrix from the corresponding uniform distributions (see Table S3).
5. Calculate all abundances ($F_{iz}$, $F_{ij}$) and frequencies of all different host types in the population ($f_{iz}$, $f_{ij}$, $\tilde{f}_{ij}$) for a population of size $N_H = 100,000$ hosts and a given value of $\phi$.
6. Based on these frequencies calculate the value of each index using data from the entire population.
7. Draw a sample $Binom(n_H, \tilde{f}_{1z})$ from the **non-infected** part of the population
8. Draw a sample from $Multi(n_I; \tilde{f}_{11}, \tilde{f}_{12}, \tilde{f}_{21}, \tilde{f}_{22})$ from the **infected part** of the population.
9. Check if all host and pathogen alleles are present in the infected sample $n_I$. If not discard the entire simulation and proceed to next simulation.
10. Calculate the indices for the sample

This procedure was repeated until either $m = 50,000$ simulations were generated for the given matrix or if a maximum number of $m_I$ attempts to generate simulations was reached.

We generated simulations for all six matrices in Table S2 for all pairwise combinations of $\phi \in \{0.05, 0.5, 0.95\}$ and $\delta \in \{0.1, 0.2, 0.3\}$ resulting in a total of 3x3=9 simulation sets with 50,000 simulations each. For all main results we fixed our sample sizes to $n_H = 1006$ and $n_I = 902$ haploid hosts.

To gain additional insights in fixing sample sizes for infected and uninfected hosts, we repeated the recipe from above for the same simulation grid for (i) $n_H = 1813$ and $n_I = 95$ and for (ii) $n_H = 95$ and $n_I = 1813$.

#### Leave-one-out cross validation for model selection with ABC

We first tested the suitability of Approximate Bayesian Computation (ABC) model choice to distinguish the six different matrices for a given combination of $\phi$ and $\delta$ by using our four indices summary statistics in an ABC. Therefore, we run a leave-one-out cross-validation for a cross-validation sample of size 500 using the function `cv4postpr` from the R-package abc [6] using the rejection algorithm and keeping the $5\%$ best simulations for each cross-validation sample. We run the cross-validation separately for the entire population and for the samples taken from the population.

#### Model choice for the top association candidates

We selected the associations with the 200 highest values for each of our indices from the human/HCV dataset. For each of these 800 associations we run ABC model choice using our simulated dataset for the different matrices for $\phi = 0.05$, $\delta = 0.1$, $n_H = 1006$ and $n_I = 902$ and our four indices as summary statistics. We run the abc model choice using the function `postpr(...,tol=0.05, method="rejection")` from the R-package `abc` v2.2.1 [6]. For each candidate association, we filtered for the model (matrix) with the highest overall Bayes factor and investigated its Bayes factors against any other model. We categorized the results as follows:

| Category | Description |
|---|---|
| Category 1 | the model with the highest Bayes factor is a non-neutral model/matrix, and the Bayes factor compared to the neutral matrix is larger than 2. |
| Category 2 | the model with the highest Bayes factor is a non-neutral model/matrix, but the Bayes factor compared to the neutral model is smaller or equal to 2. |
| Category 3 | the model with the highest Bayes factor is the neutral model. |

We considered all associations falling into Category 1 as confident assignments to a non-neutral matrix. In addition we summarized the model choice results for all 200 top associations of a given index (highest value) by counting the number of times each best matrix was the single best matrix (all Bayes factors compared to any other model $> 2$) or it was competing with any combination of other matrices (all other matrices with Bayes factor

$< 2$).

Table S2 Valid configuration of the extreme matrices.

| **neutral infection matrix** $\mathcal{A}_N$ | **differential host resistance** $\mathcal{A}_H$ | **differential pathogen infectivity** $\mathcal{A}_P$ | **matching alleles (MA)** $\mathcal{A}_{\mathcal{MA}}$ | **gene-for-gene (GFG)** $\mathcal{A}_{\mathcal{GFG}}$ | **inverse gene-for-gene** $\mathcal{A}_{i\mathcal{GFG}}$ |
|---|---|---|---|---|---|
| $\begin{pmatrix} 1 & 1 \\ 1 & 1 \end{pmatrix}$ | $\begin{pmatrix} 1 & 1 \\ 0 & 0 \end{pmatrix}$ $\begin{pmatrix} 0 & 0 \\ 1 & 1 \end{pmatrix}$ | $\begin{pmatrix} 1 & 0 \\ 1 & 0 \end{pmatrix}$ $\begin{pmatrix} 0 & 1 \\ 0 & 1 \end{pmatrix}$ | $\begin{pmatrix} 1 & 0 \\ 0 & 1 \end{pmatrix}$ $\begin{pmatrix} 0 & 1 \\ 1 & 0 \end{pmatrix}$ | $\begin{pmatrix} 0 & 1 \\ 1 & 1 \end{pmatrix}$ $\begin{pmatrix} 1 & 0 \\ 1 & 1 \end{pmatrix}$ $\begin{pmatrix} 1 & 1 \\ 0 & 1 \end{pmatrix}$ $\begin{pmatrix} 1 & 1 \\ 1 & 0 \end{pmatrix}$ | $\begin{pmatrix} 1 & 0 \\ 0 & 0 \end{pmatrix}$ $\begin{pmatrix} 0 & 1 \\ 0 & 0 \end{pmatrix}$ $\begin{pmatrix} 0 & 0 \\ 1 & 0 \end{pmatrix}$ $\begin{pmatrix} 0 & 0 \\ 0 & 1 \end{pmatrix}$ |

Table S3 Scheme for randomly simulating deviations from the extreme matrixes.

| **neutral infection matrix** $\mathcal{A}_N$ | **differential host resistance** $\mathcal{A}_H$ | **differential pathogen infectivity** $\mathcal{A}_P$ | **matching alleles (MA)** $\mathcal{A}_{\mathcal{MA}}$ | **gene-for-gene (GFG)** $\mathcal{A}_{\mathcal{GFG}}$ | **inverse gene-for-gene** $\mathcal{A}_{i\mathcal{GFG}}$ |
|---|---|---|---|---|---|
| $\begin{pmatrix} \mathcal{U}_1 & \mathcal{U}_1 \\ \mathcal{U}_1 & \mathcal{U}_1 \end{pmatrix}$ | $\begin{pmatrix} \mathcal{U}_1 & \mathcal{U}_1 \\ \mathcal{U}_0 & \mathcal{U}_0 \end{pmatrix}$ $\begin{pmatrix} \mathcal{U}_0 & \mathcal{U}_0 \\ \mathcal{U}_1 & \mathcal{U}_1 \end{pmatrix}$ | $\begin{pmatrix} \mathcal{U}_1 & \mathcal{U}_0 \\ \mathcal{U}_1 & \mathcal{U}_0 \end{pmatrix}$ $\begin{pmatrix} \mathcal{U}_0 & \mathcal{U}_1 \\ \mathcal{U}_0 & \mathcal{U}_1 \end{pmatrix}$ | $\begin{pmatrix} \mathcal{U}_1 & \mathcal{U}_0 \\ \mathcal{U}_0 & \mathcal{U}_1 \end{pmatrix}$ $\begin{pmatrix} \mathcal{U}_0 & \mathcal{U}_1 \\ \mathcal{U}_1 & \mathcal{U}_0 \end{pmatrix}$ | $\begin{pmatrix} \mathcal{U}_0 & \mathcal{U}_1 \\ \mathcal{U}_1 & \mathcal{U}_1 \end{pmatrix}$ $\begin{pmatrix} \mathcal{U}_1 & \mathcal{U}_0 \\ \mathcal{U}_1 & \mathcal{U}_1 \end{pmatrix}$ $\begin{pmatrix} \mathcal{U}_1 & \mathcal{U}_1 \\ \mathcal{U}_0 & \mathcal{U}_1 \end{pmatrix}$ $\begin{pmatrix} \mathcal{U}_1 & \mathcal{U}_1 \\ \mathcal{U}_1 & \mathcal{U}_0 \end{pmatrix}$ | $\begin{pmatrix} \mathcal{U}_1 & \mathcal{U}_0 \\ \mathcal{U}_0 & \mathcal{U}_0 \end{pmatrix}$ $\begin{pmatrix} \mathcal{U}_0 & \mathcal{U}_1 \\ \mathcal{U}_0 & \mathcal{U}_0 \end{pmatrix}$ $\begin{pmatrix} \mathcal{U}_0 & \mathcal{U}_0 \\ \mathcal{U}_1 & \mathcal{U}_0 \end{pmatrix}$ $\begin{pmatrix} \mathcal{U}_0 & \mathcal{U}_0 \\ \mathcal{U}_0 & \mathcal{U}_1 \end{pmatrix}$ |

where $\mathcal{U}_0 \sim \mathcal{U}(0, \delta), \mathcal{U}_1 \sim \mathcal{U}(1-\delta, 1)$ and $\delta \in [0, 1]$.

### Genome data analysis

#### Inference of past population dynamics in HCV

We examined the past HCV population size change based on the European individuals using a Coalescent Bayesian Skyline model, which extends the basic coalescent model by allowing the population size to vary over time, and estimates the changes in effective population size through time from SNP data [8]. We analyzed the HCV nucleotide alignment with a Bayesian Skyline Plot coalescent prior in BEAST v2.4 [3, 8]. Bayesian analyses for each transmission clade employ the GTR model of nucleotide substitution using a strict clock model with a clock rate of $0.79e{-}3$ substitutions per site per y (s/s/y) ([14, 15, 17]). The GTR model divides the time between present and the root of the tree (the tMRCA) into segments and then estimates the effective population size ($N_e$) for each segment ([17]). The present time point in the analysis was placed to the time of data sampling (2015)([10]). We used the remaining prior distribution of the Bayesian skyline population parameter sizes with default setting ([17]). One MCMC run was conducted in the analyses, with 10 million generations and the trees were sampled every 1000 generations, with the first 10% discarded as burn-in.

#### Pre-processing of the human data

For the infected human data, we obtained human genotype data from 583 patients enrolled in the BOSON study, a phase 3 randomized open-label trial, which evaluated the safety and effectiveness of Sofosbuvir (Foster et al., 2015). We downloaded genotype data of 541 patients, generated by an Affymetrix UK Biobank array, from the European Genome-phenome Archive (accession: EGAS00001002324, [2]). This array genotypes over 800,000 genome-wide single-nucleotide polymorphisms (SNPs) and covers variants in the HLA region, which is known to be important in human immune response [2]. The available human genotype data was matched to the available HCV whole-genome sequences from GenBank (accessions: KY620313–KY620880) and resulted in 541 human-virus pairs of mainly self-reported European (451 individuals) and Asian (74 individuals) ancestry. Quality control and filtering consisted of two steps: (i) identification of SNPs showing a significant deviation from Hardy-Weinberg equilibrium (HWE) using PLINK2 (`www.cog-genomics.org/plink/2.0/`)

([5]) with $P < 1e{-6}$ (Literature reported significance thresholds for HWE are between 0.001 and $5.7e{-7}$) ([4, 13]) and (ii) the removal of all markers with minor allele frequency (MAF) $P < 0.2$ (using PLINK2). Further, we pruned for LD within a 50 kilobase (Kb) window, using a step size of 10 and an $r^2$ threshold of 0.1. The resulting pruned datasets were subjected to principal component analysis (PCA) using PLINK2 ([5]) to evaluate the underlying population stratification. The PCA results were visualized in R, version 4.0.2. In order to avoid potential confounding effects of population stratification, we opted to limit our analysis to the European dataset exclusively. After quality control and filtering of the human genotype data, 333,655 SNPs (326,520 SNPs for infected European dataset) were available for further analysis.

For the non-infected sample, 1000 Genomes phase III data (1000GenomesIII) were downloaded as processed VCF files from the International Genome Sample Resource (IGSR) website that maintains access to 1000 Genomes Project data [1, 9]. The 2504 unrelated samples were concatenated into a multi sample and multi chromosome VCF file using vcftools ([7]). Further, we included only individuals from European countries and we matched the variants to all SNPs of the infected European human sub-sample (from [2]), in order to obtain all 326,520 SNPs (see pre-processing of infected sub-sample).

#### Pre-processing of virus data

We downloaded the viral data (nucleotide and protein) [2] from NCBI GenBank (accessions KY620313–KY620880). Following [2], we generated whole-genome viral consensus sequences (nucleotide and protein) for each patient using MAFFT (v.7.429) [11]. Based on the protein consensus sequences we inferred the phylogenetic relationship using RaxML v.8.2.4 (rate distribution model PROTGAMMAWAG, 100 bootstraps) ([18]). Next, we analysed viral population structure based on the viral nucleotide consensus sequences. Therefore, we first trimmed the multisequence alignments using the function msaTrim (gap.end=0.04, gap.mid=0.08) of the R-package microseq (v.2.1.5) ([16]). Missing values were inputed based on the sequence with the smallest overall distance by using the function dist.dna (model=K80) (R-package ape [12]) and customized R-codes. On these data we run a principle component analysis using the prcomp from the R-stats package (v.3.6.2) (R Core Team).

To prepare the data for plink we first filtered for all amino acids where the missing value rate was $< 1.452e{-3}$ ($< 1.437e{-3}$ for European dataset). Then we filled all missing values with "-" and coerced low frequency amino acids (n <3) into the next higher amino acid frequency group. Amino acid positions with more than two amino acids present were converted into binary response variables by splitting the original column into several columns. In each column one of the amino acids was then coded as 1 and all the other amino acids as zero. In example if there is a column where: I=443 (Leucine), T=50 (Threonine), V=38 (Valine), S=5 (Serine), X=4 (Glutamine) an F=1 (Phenylalanine), we would first add the F=1 amino acid to the X (Glutamine) group. Hence, we end up with four amino acid categories with the following (allele) frequencies f(I)= 0.818854, f(T)=0.092421, f(V)=0.070240, f(S)=0.009242, f(X+F)=0.009242). There are three amino acid categories with an allele frequency $> 0.02$ and accordingly the column is converted into three binary columns. In the first of these columns all samples with a Leucine (I) at this position would be encoded by 1 and all other samples with a 0. In the second column, all individuals with a Tyrosine (T) would be encoded by a 1 and all other individuals with a 0 and in the third column all individuals with a Valine (V) are encoded by a 1 and all other individuals with a zero. This resulted in a total of 1,779 columns (1,626 columns for European sub-sample) of the virus alignment, split into 2,815 columns (2,751 for European sub-sample) after binary conversion, that were used as input for the association analysis. After that, we set a new MAF of 0.2 which resulted in 213 columns (208 columns for the European sub-sample).

[3] R. Bouckaert, T. G. Vaughan, J. Barido-Sottani, S. Duchêne, M. Fourment, A. Gavryushkina, J. Heled, G. Jones, D. Kühnert, N. De Maio, et al. Beast 2.5: An

advanced software platform for bayesian evolutionary analysis. *PLoS computational biology*, 15(4):e1006650, 2019.

[10] G. R. Foster, N. Afdhal, S. K. Roberts, N. Bräu, E. J. Gane, S. Pianko, E. Lawitz, A. Thompson, M. L. Shiffman, C. Cooper, W. J. Towner, B. Conway, P. Ruane,

M. Bourlière, T. Asselah, T. Berg, S. Zeuzem, W. Rosenberg, K. Agarwal, C. A. Stedman, H. Mo, H. Dvory-Sobol, L. Han, J. Wang, J. McNally, A. Osinusi, D. M. Brainard, J. G. McHutchison, F. Mazzotta, T. T. Tran, S. C. Gordon, K. Patel, N. Reau, A. Mangia, and M. Sulkowski. Sofosbuvir and velpatasvir for hcv genotype 2 and 3 infection. *New England Journal of Medicine*, 373(27):2608–2617, 2015. doi: 10.1056/NEJMoa1512612. URL https://doi.org/10.1056/NEJMoa1512612. PMID: 26575258.
